# Caveolin-1 temporal modulation enhances trastuzumab and trastuzumab-drug conjugate efficacy in heterogeneous gastric cancer

**DOI:** 10.1101/2021.07.13.452260

**Authors:** Patrícia M. R. Pereira, Komal Mandleywala, Sébastien Monette, Melissa Lumish, Kathryn M. Tully, Mike Cornejo, Audrey Mauguen, Ashwin Ragupathi, Marissa Mattar, Yelena Y. Janjigian, Jason S. Lewis

## Abstract

Resistance mechanisms and heterogeneity in HER2-positive gastric cancers (GC) limit trastuzumab benefit in 32% of patients, and other targeted therapies have failed in clinical trials. Using genomic data from patient tissue, patient-derived xenografts (PDXs), partially humanized biological models, and HER2-targeted imaging we identified caveolin-1 (CAV1) as a complementary biomarker in GC selection for trastuzumab therapy. In retrospective analyses of samples from patients enrolled on trastuzumab trials, the CAV1-high profile was associated with low membrane HER2 density and reduced patient survival. We found a negative correlation between CAV1 tumoral protein levels – a major protein of cholesterol-rich membrane domains – and trastuzumab-drug conjugate TDM1 tumor uptake. Finally, CAV1 depletion using knockdown or pharmacologic approaches was shown to increase HER2-directed immunoPET uptake and TDM1 efficacy in GC with incomplete HER2 membranous reactivity. In support of these findings, background statin use in patients is associated with enhanced antibody efficacy. Together, this work provides mechanistic justification and clinical evidence that require prospective investigation of HER2-targeted therapies combined with statins to delay drug resistance in GC.

**SIGNIFICANCE:** This study identifies how CAV1 protein expression and statin use relate to GC response in HER2-targeted imaging and therapeutic approaches. In addition, to support the synergy of CAV1 depletion with TDM1 observed in mouse models, we demonstrate that statin users had better clinical responses to antibody-based therapies in HER2^+^ GC.

## INTRODUCTION

Human epidermal growth factor receptor 2 (HER2) alterations, including overexpression, amplification, and other mutations, occur in breast and gastric tumors [1, 2]. Although several targeting agents are effective in treating HER2-positive breast tumors [1, 2], not all HER2-expressing tumors benefit from HER2-targeted therapies (reviewed in [2]). Beyond trastuzumab [3] and trastuzumab deruxtecan [4], clinical trials have failed to demonstrate efficacy of other HER2-targeted therapies (pertuzumab, TDM1) in the first and later treatment lines for gastric cancer (GC) [5, 6].

HER2 belongs to the EGFR family of receptor tyrosine kinases (RTKs). HER2 overexpression and/or amplification exists in approximately 15–20% of breast or gastric cancers [2]. Receptor dimerization often occurs in HER2-overexpressed tumors, switching on the kinase domains that trigger signaling pathways [1]. The anti-HER2 antibody trastuzumab is the standard-of-care treatment for metastatic and early-stage HER2-positive breast cancer [2] and first-line therapy in combination with chemotherapy for GC [3]. Mechanistically, trastuzumab hinders receptor dimerization, engenders receptor internalization and degradation, inhibits the PI3K-AKT signaling pathway, and stimulates antibody-dependent cytotoxicity (ADCC). The natural killer (NK) cells are notable players in trastuzumab-mediated ADCC: antibody efficacy correlates preclinically and clinically with NK cell function and stimulation [7, 8], and a loss in activity occurs in Fcγ-RIIIA knockout mice [9]. Other approved HER2-targeting agents for breast cancer include the antibody pertuzumab, tyrosine kinase inhibitors, and antibody-drug conjugates (ADCs). HER2-targeted therapies are also being tested in the biliary tract, colorectal, non-small-cell-lung and bladder cancers, owing to HER2 overexpression in these solid tumors [2].

In GC, intrinsic or acquired resistance to trastuzumab results in inevitable disease progression. Furthermore, trastuzumab conjugated with the tubulin-binding agent DM1 (ado-trastuzumab emtansine, TDM-1), and anti-HER2 antibody combinations (trastuzumab and pertuzumab) plus chemotherapy have all failed to improve overall survival in clinical trials for patients with GC [5, 6]. Heterogeneous expression of HER2 in GC [2, 10–12] is one reason why the pertuzumab and TDM1 failed in clinical trials of GC [5, 6]. Notably, the selection of patients for HER2-targeted therapies based on the assessment of receptor status through immunohistochemistry (IHC) and *in situ* hybridization (ISH) of tumor biopsy specimens incompletely characterizes the cellular dynamics of HER2 and the heterogeneity of its expression in GC [13, 14]. Complementary techniques and biomarkers are therefore needed during tumor selection and therapeutic response monitoring. These include DNA sequencing of tumor and cell-free DNA as well as whole-body HER2-directed functional positron emission tomography (PET). Zirconium-89 (^89^Zr)-labeled anti-HER2 antibody immunoPET has shown potential as a complementary technique to image GC heterogeneity, which may be predictive of subsequent drug response within weeks of therapy initiation [15, 16].

Receptor endocytosis and recycling processes contribute to HER2 heterogeneity and membrane dynamics [17], affecting antibody–tumor binding and subsequent efficacy and ADCC-mediated mechanisms [18, 19]. Although there is evidence for HER2 localization in cholesterol-rich structures — caveolae — at the cell surface [20], studies have also shown occasional HER2 presence in clathrin-coated pits [17] or endophilin-mediated mechanisms [21]. Caveolin-1 (CAV1), the major structural protein of caveolae, negatively correlates with membrane HER2 and affects trastuzumab-tumor binding [22–28]. Endocytic trafficking systems also influence TDM1 efficacy [19]. While CAV1-dependent endocytosis enhances cancer cells’ chemosensitivity to TDM1 [29], others have shown a role for caveolae-mediated endocytosis in TDM1 resistance [25, 28]. Because cholesterol is essential for caveolae assembly and stability [30], cholesterol-depleting drugs are used in preclinical studies to modulate CAV1 protein levels [31–35]. In this context, statins are promising pharmacologic CAV1 modulators, since they are FDA-approved drugs and prescribed to millions of people worldwide for the treatment of hypercholesterolemia [31]. In preclinical models, statins enhance HER2 confinement at the cell surface [22, 36], increase HER2-directed immunoPET uptake and enhance trastuzumab systemic efficacy in GC with non-predominant HER2 membrane staining [22].

In this work, we retrospectively validate CAV1 as a complementary biomarker for the selection of patients with GC for HER2-targeted therapies. Tumors with high CAV1 correspond with low HER2 density at the cell surface and, in trastuzumab trials, to patients with low survival rates. Using GC cell lines and PDXs with varying levels of CAV1, we show that TDM1, an ADC that targets HER2, combined with lovastatin, a small molecule that depletes cholesterol in ways that modulate CAV1 protein expression, improves immunoPET and response rates better than either does alone in heterogeneous GC. Mechanistically, statins enhance the disruption of downstream signaling and NK-mediated ADCC. Importantly, we validated these preclinical findings in retrospective analyses of patient-level data from clinical studies of HER2-targeted therapies in GC patients.

## RESULTS

### CAV1-high and CAV1-low gastric tumors have distinct HER2-signaling profiles

Previous studies of HER2-positive tumor models implied a role for CAV1 in trastuzumab or TDM1 binding and efficacy [22, 24, 25, 28, 29]. These findings provided the rationale for a retrospective study to identify genetic features of CAV1-low and CAV1-high HER2+ GC. Eligible samples were obtained from previous trials at Memorial Sloan-Kettering Cancer Center (MSK) of HER2+ GC patients treated with trastuzumab. Samples were from predominantly male patients (80% male versus 20% female) with a median age of 61 years (range 28–87). HER2-positivity was defined as IHC 3+, IHC 2+ and HER2:CEP17 FISH ratio ≥ 2.0, or *ERBB2* amplification by next-generation sequencing. **Table 1** summarizes patient characteristics. The cohort consists of 46 patients with stage IV (74 %), stage III (17%), or stage II (9%) HER2+ GC disease at the time of diagnosis. All patients were stage IV at the point when trastuzumab therapy was initiated. Samples obtained from patients enrolled on trastuzumab trials (9/46 tumor samples were from patients that received other therapies prior to trastuzumab) were analyzed for CAV1 IHC and genomic alterations (**Fig. 1A,B**). This cohort was comprised of samples from primary tumors (43%) or metastases (57%). CAV1 IHC optimization used tissues with varying levels of CAV1 (**Supplementary Figs. 1,2**). CAV1 staining at the membrane of GC was classified as 0/1+ CAV1-low and 2+/3+ CAV1-high (**Fig. 1A**). CAV1-high and CAV1-low IHC were detected respectively in 26% and 74% of HER2+ GC.

**Figure 1.**
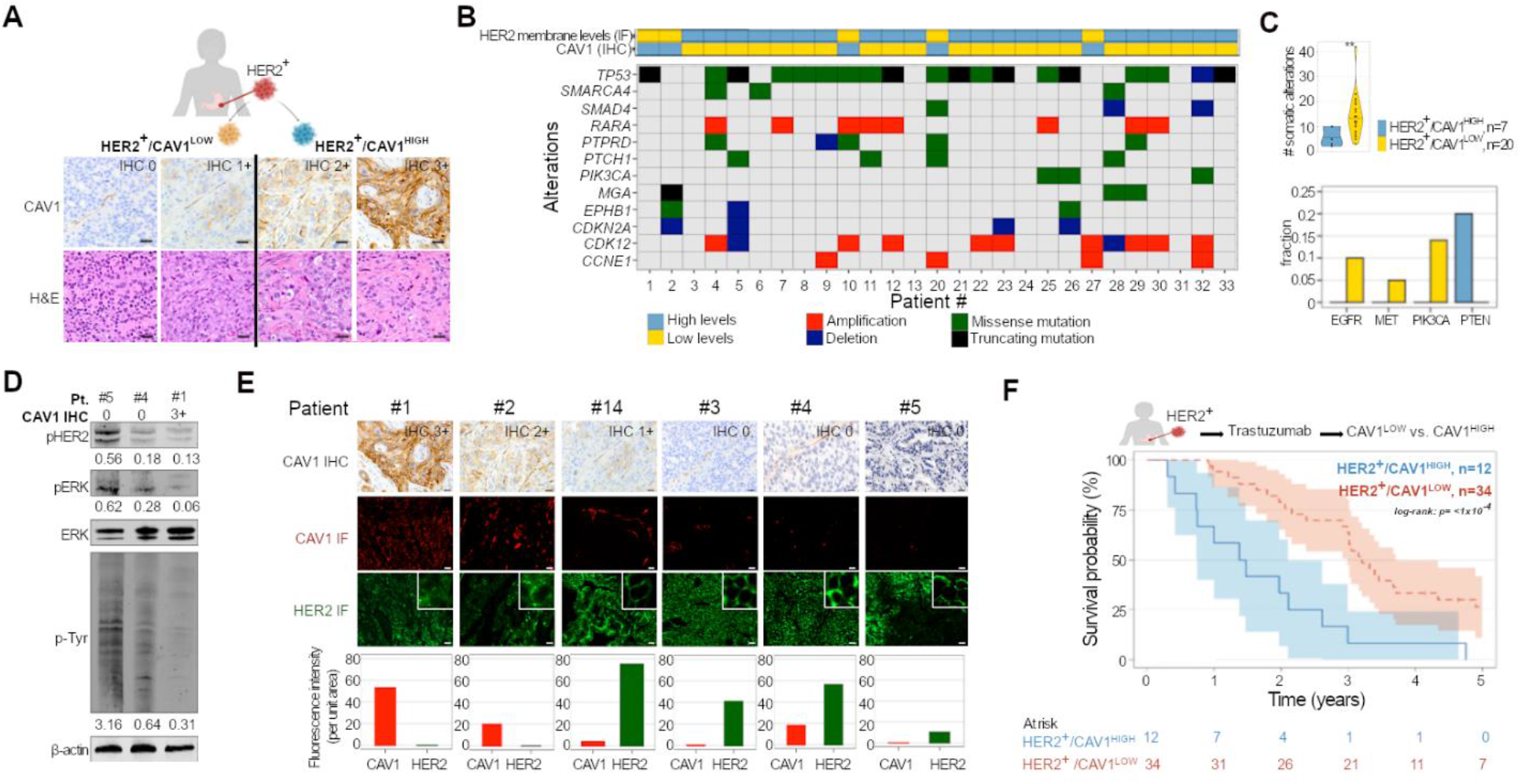
HER2 membrane levels and trastuzumab efficacy depend on CAV1 protein levels. (**A**) Immunohistochemical detection and scoring intensity of CAV1 in HER2-expressing gastric tumor tissues. Only CAV1 reactivity at the cell membrane of tumor cells was considered for scoring. CAV1 0/1+: CAV1-low. CAV1 2+/3+: CAV1-high. (**B**) The 12 most predominant genomic alterations present in CAV1-low versus CAV1-high HER2^+^ gastric tumor tissues. HER2 membrane levels are classified as high versus low based on quantification of IF staining shown in **Supplementary Fig. 2**. Patient 1 to Patient 33 are IDs for all HER2^+^ gastric tumor tissues analyzed in the study. (**C**) Number of genomic alterations, *EGFR* and *MET* amplification, *PIK3CA* and *PTEN* mutations present in HER2^+^CAV1^LOW^ versus HER2^+^CAV1^HIGH^ gastric tumor tissues. HER2^+^CAV1^LOW^ (*n* = 20), HER2^+^/CAV1^HIGH^ (*n* = 7). (**D**) Western blot analyses of HER2, ERK, and Tyr phosphorylation in HER2^+^/CAV1^LOW^ (patient #4 and patient #5) and HER2^+^/CAV1^HIGH^ (patient #1) gastric tumor tissues. (**E**) CAV1 IHC, confocal images, and quantification of immunofluorescence staining of HER2 (green color) and CAV1 (red color) in human HER2-expressing gastric tumors. CAV1-high: CAV1 2+/3+ (patient #1 and patient #2). CAV1-low: CAV1 1+/0 (patient #14 and patients #3–5). The graphs plot protein fluorescence intensity per unit area, calculated by quantifying IF images (mean ± S.E.M, *n* = 3). Scale bar, 50 μm. (**F**) Kaplan-Meier analyses of CAV1 expression and GC disease outcome in patients treated with trastuzumab. Patients with HER2^+^/CAV1^HIGH^ (blue color, *n* = 12 patients) phenotype have a worse survival than HER2^+^/CAV1^LOW^ (red, *n* = 34 patients). Log rank; p < 1 × 10^-4^.

**Table 1.** Patient characteristics.

| **Characteristics** | **N (%)** |
| --- | --- |
| Total patients | 46 |
| Sex, median |  |
| Male | 37 (80) |
| Female | 9 (20) |
| Age at diagnosis, median | 61 (between 28 to 87) |
| Therapy Prior to Trastuzumab | 9 (19.5) |
| EOX | 2 |
| Carboplatin/paclitaxel/RT | 2 |
| Carboplatin/taxol/RT | 1 |
| Modified DCF | 1 |
| FOLFOX | 2 |
| FOLFOX/regorafenib 2. 5-FU/regorafenib | 1 |
| Stage of disease at the time of diagnosis |  |
| Stage IV | 34 (74) |
| Stage III | 8 (17) |
| Stage II | 4 (9) |
| Stage of disease at the time of initiating trastuzumab |  |
| Stage IV | 46 (100) |
| HER2 IHC at diagnosis |  |
| 3+ | 36 (78) |
| 2+ | 10 (22) |
| HER2-positivity retained after initiating trastuzumab | 14 (30) |
| Sample type |  |
| Primary | 20 (43) |
| Metastasis | 26 (57) |
| Sample location |  |
| Liver | 12 (26) |
| Stomach | 10 (22) |
| Esophagus | 6 (13) |
| Esophagogastric junction | 6 (13) |
| Lung | 5 (11) |
| Brain | 4 (8.7) |
| Scalp | 1 (2.1) |
| Skin | 1 (2.1) |
| Peritoneum | 1 (2.1) |
| **CAV1 IHC** |  |
| HER2<sup>+</sup>/CAV1<sup>HIGH</sup> tumors | 12 (26) |
| HER2<sup>+</sup>/CAV1<sup>LOW</sup> tumors | 34 (74) |
| **Statin use** | 19 (41) |
| Statin use prior trastuzumab therapy | 12 (26) |
| Prior trastuzumab at PDX biopsy |  |
| Yes | 36 (78) |
| No | 10 (22) |
| Stage of disease at the time of PDX biopsy |  |
| Stage IV | 40 (87) |
| Stage III | 6 (13) |
| EOX: chemotherapy combination of epirubicin, oxaliplatin, and capecitabine (Xeloda).<br>RT: radiation therapy<br>Modified DCF: modified docetaxel, cisplatin, fluorouracil |  |

At the time of the retrospective study, MSK-IMPACT data was available for 33 of 46 HER2^+^ GC samples. This methodology uses a hybridization-based exon capture design to detect somatic single-nucleotide variants, small insertions and deletions, copy-number alterations, and structural rearrangements [15, 37]. We, therefore, used MSK-IMPACT data to discover somatic alterations in HER2^+^/CAV1^LOW^ and HER2^+^/CAV1^HIGH^ GC, (**Supplementary Fig. 3**). The most frequently altered genes in a cohort of 27 samples studied were *TP53*, *SMARCA4*, *RARA*, *PTPRD*, *PTCH1*, *PIK3CA*, *MGA*, *EPHB1*, *CDKN2A*, *CDK12*, and *CCNE1* (**Fig. 1B**). Overall, CAV1-high GC exhibited a relatively low number of mutations compared with CAV1-low tumors (**Fig. 1C, Supplementary Fig. 3**). *EGFR*- and *MET*-amplification and *PIK3CA* mutations associated with low *PTEN* levels were observed respectively in 10%, 5%, and 15% of HER2^+^/CAV1^LOW^ cases but absent in HER2^+^/CAV1^HIGH^ tumors (**Fig. 1C, Supplementary Fig. 3**). CAV1-low tumors were also associated with higher activation of HER2, ERK, and Tyr than CAV1-high tumors (**Fig. 1D**). Taken together, our results suggest different signaling profiles in CAV1-high versus CAV1-low GC.

### CAV1-low profile predicts favorable GC response in patients undergoing trastuzumab therapy

Immunofluorescence staining of patient samples revealed high membrane density of HER2 receptors in CAV1-low GC (**Fig. 1B, Fig. 1E, Supplementary Fig. 2**). Conversely, non-homogeneous HER2 membrane staining prevailed in CAV1-high tumors. Western blot studies of a panel of 6 GC cell lines supported the inverse correlation between HER2 and CAV1 protein expression (**Supplementary Fig. 4A**). These observations are consistent with previous results in preclinical models of HER2-expressing tumors [22–25]. We then sought to determine if tumoral CAV1 was associated with survival in patients undergoing HER2-targeted therapy. We compared patient survival during trastuzumab therapy in stratified CAV1-high and CAV1-low GC. These retrospective analyses used information about the 46 patients described above to determine the association of CAV1 IHC and HER2 membrane staining with trastuzumab responses and overall survival in trials of HER2+ GC (**Table 1**). In retrospective analyses performed in this study, the CAV1-low profile (34 of 46) corresponds to tumors with homogeneous surface receptors and predicts favorable patient response to trastuzumab therapy (**Fig. 1F**).

### CAV1 depletion increases TDM1 accumulation in GC with incomplete patterns of HER2 membrane staining

Membrane-localized receptors and trafficking are important in the therapeutic efficiency of ADCs [19]. We next hypothesized that HER2-positive CAV1-high and CAV1-low GC have different susceptibility to ADCs based on different HER2 membrane availability and internalization rates. To test whether CAV1 protein levels influence TDM1 binding and trafficking, we first used a panel of GC cell lines (**Supplementary Fig. 4A,B**): a nontransformed HER2+ GC cell line (WT, NCIN87) and three GC cell lines (AGS, KATOIII, and SNU1) stably expressing HER2 (LV-HER2). In addition to the generation of KATOIII, AGS, and SNU1 sublines stably expressing HER2, we also attempted to express HER2 in MKN45 and SNU5 GC cells. Interestingly, the protocols herein used did not allow for the successful generation of LV-HER2 in cell lines containing the highest CAV1 expression (**Supplementary Fig. 4B)**. While CAV1 protein negatively correlates with membrane HER2 [22], CAV1 role in HER2 transduction processes are less clear. NCIN87 WT, AGS LV-HER2, KATOIII LV-HER2, and SNU1 LV-HER2 were used for quantification of TDM1 binding or internalization (**Fig. 2A**). ADC binding to GC cells, as determined using ^89^Zr-labeled TDM1, negatively correlates with HER2 total levels (Spearman’s rank correlation coefficient (*r*) of −0.92, *p*=0.0012) and positively correlates with HER2-to-CAV1 ratios (*r*=0.89, *p*=0.00338). We next determined if differences in TDM1 binding would affect ADC internalization in GC cells. ADC internalization was measured using TDM1 conjugated with the pH-sensitive dye (pHrodo-TDM1) that only fluoresces in acidic environments, such as the lysosome [19]. We detected the highest pHrodo-TDM1 signals in GC cells containing the highest HER2-to-CAV1 protein ratios (**Fig. 2A**). Similarly, others have recently described TDM1 uptake and endocytosis regardless of the total protein levels of HER2 [19, 25]. Altogether, these findings support a positive correlation between HER2-to-CAV1 ratios and TDM1 binding and internalization in GC cells.

**Figure 2.**
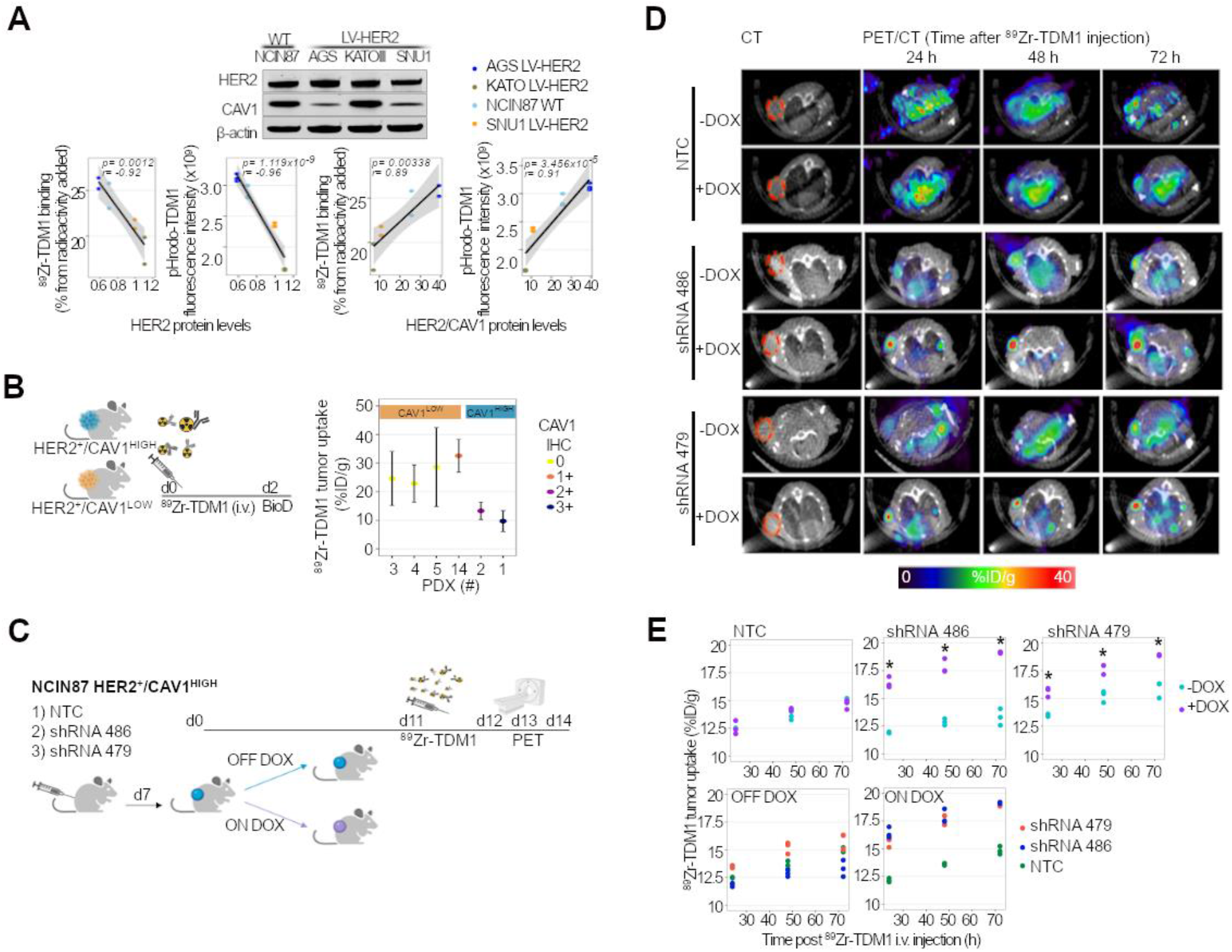
TDM1-tumor binding and internalization depend on CAV1 protein levels. (**A**) TDM1 binding and internalization in NCIN87 GC cells wild-type (WT) and AGS, KATOIII, SNU1 GC sublines stably expressing HER2 (LV-HER2). For the binding assays, cells were incubated with ^89^Zr-labeled TDM1 (1 μCi, 0.25 μg) for 1 h at 4°C. TDM1 internalization was performed by incubating GC cells with 1 μg/mL of pHrodo-TDM1 for 30 min at 4°C, and then releasing at 37°C between 0.5 to 48 h. The levels of ^89^Zr-TDM1 binding or pHrodo-TDM1 internalization (*y*-axis) were plotted against HER2 or ratio of HER2-to-CAV1 protein levels (*x*-axis). The non-parametric one-tailed Spearman test was used to determine the correlation coefficients. HER2 or HER2/CAV1 protein levels were determined by western blot analysis from total extracts (upper panel). (**B**) [^89^Zr]Zr-DFO-TDM1 uptake in HER2-expressing gastric PDXs containing varying levels of CAV1. NSG mice bearing subcutaneous PDXs were intravenously administered with [^89^Zr]Zr-DFO-TDM1 (6.66–7.4 Mbq, 45–50 μg protein) and biodistribution performed at 48 h p.i. of ^89^Zr-labeled antibody. PDX IDs in this figure match patient IDs shown in **Figure 1**. Points, *n* = 5 mice per group, mean ± S.E.M. %ID/g, percentage of injected dose per gram. (**C-E**) Athymic nude mice bearing s.c. NCIN87 shRNA NTC, shRNA 486, or shRNA 479 xenografts were orally administered with 10 mg/mL of Dox (ON DOX) or PBS (OFF DOX) for 11 days. On day 11, mice were intravenously administered with [^89^Zr]Zr-DFO-TDM1 (6.66–7.4 Mbq, 45–50 μg protein). PET images (**D**) were recorded at 24, 48, and 72 h p.i. [^89^Zr]Zr-DFO-TDM1. The percentage of injected dose per gram (%ID/g) of TDM1 in tumors (**E**) was calculated by quantifying regions of interest (ROIs) in the PET images. \**P < 0.05* based on a Student’s *t*-test, *n* = 3.

Preclinical imaging studies were then performed using HER2^+^ gastric PDXs (78% of PDXs were obtained from patients prior initiating trastuzumab; **Table 1**) to establish the significance of CAV1 levels on TDM1-tumor binding. The PDX tissues were confirmed to match the parent tissue shown in **Fig. 1** by MSK-IMPACT data. Examination of H&E and IHC stained sections excluded the possible presence of B cell lymphomas in PDXs associated with Epstein-Barr Virus (EPV) [38, 39] (**Supplementary Fig. 4C**; carcinomas: pancytokeratin^+^/CD45^-^/CD20^-^, lymphomas: pancytokeratin^-^/CD45^+^/CD20^+^). PDXs containing lymphoma were excluded from preclinical studies (*n*=13). At 48 h post-injection of ^89^Zr-labeled TDM1, CAV1-low PDXs had uptakes ranging from 22.8 ± 6.5 to 32.5 ± 5.7 percentage of injected dose per gram of PDX (%ID/g), while CAV1-high PDXs yielded uptakes ranging from 9.7 ± 3.6 to 13.3 ± 3.0 %ID/g (**Fig. 2B**). The lower TDM1 accumulation in CAV1-high xenografts, when compared with CAV1-low tumors, prompted us to interrogate if *in vivo* genetic depletion of CAV1 would boost ADC uptake. To this end, we used CAV1-high NCIN87 GC cells containing incomplete HER2 surface density [22] to develop a Tet-On system of CAV1 knockdown in the presence of doxycycline (Dox); **Supplementary Fig. 5A,B**. We performed *in vivo* studies in mice bearing subcutaneous (s.c.) NCIN87 shRNA 486 or shRNA 479 xenografts. Control experiments included non-targeting control (NTC) shRNA xenografts. Mice were orally administered saline (OFF DOX) or Dox (ON DOX) for 11 days before ^89^Zr-labeled TDM1 injection (**Fig. 2C**). Transversal PET images of the saline cohort showed a gradual accretion of immunoPET signal between 24 to 72 h into the HER2-positive tumors (**Fig. 2D**). Antibody uptake was similar in OFF DOX and ON DOX cohorts of control shRNA NTC xenografts. On the other hand, xenografts of shRNA 486 or shRNA 479 showed a remarkably higher tumor uptake in ON DOX groups when compared with OFF DOX cohorts. Quantitation of the signal in tumors’ regions of interest (ROI) further endorsed our findings from PET imaging (**Fig. 2E**). To temporally knockdown CAV1, ON/OFF Dox cohorts included mice treated with Dox for 7 days and saline for 4 days, (**Supplementary Fig. 5C,D)**. The TDM1-tumor uptake using a ON/OFF Dox schedule was comparable in mice having NCIN87 shRNA NTC, shRNA 486, or shRNA 479 xenografts (**Supplementary Fig. 5E)**. These results indicate that CAV1 knockdown enhances TDM1 binding to HER2^+^/CAV1^HIGH^ NCIN87 xenografts.

### Statin-mediated CAV1 modulation is temporal and enhances TDM1 binding

Premised on our findings using the Tet-on system (**Fig. 2D,E**), we explored *in vivo* CAV1 modulation employing an FDA-approved pharmacologic approach with potential for clinical translation. Given that the cholesterol-depleting drug lovastatin modulates CAV1 [22, 40, 41], we sought to determine whether lovastatin would enhance TDM1–tumor binding. Fluorescence microscopy demonstrated ADC localization at the cell surface and intracellular compartments of control cells (**Fig. 3A**). However, TDM1 exhibits predominant surface staining after 1.5 h incubation time with lovastatin when compared with no-statin. At 48 h after cells incubation with TDM1, the ADC shows similar localization in both control and statin groups. Using two different batches of the pHrodo-TDM1 conjugate, we consistently observed that NCIN87 control cells internalize more TDM1 than lovastatin-treated cells for the initial 2 h experiment (**Fig. 3B, Supplementary Fig. 6A**). Additional studies with 89Zr-labeled TDM1 at early time points demonstrate that lovastatin decreases ADC recycling to the cell membrane (**Fig. 3C**). At times longer than 2 h, TDM1 internalization is similar in control and lovastatin-incubated cells and co-localizes with the lysosomal-associated membrane protein 1 (LAMP-1, **Fig. 3D**). Consistently, lovastatin does not alter the ubiquitination of immunoprecipitated HER2 (**Supplementary Fig. 6B**). These data suggest that at early incubation times, lovastatin enhances TDM1 binding to the surface of GC cells and delays ADC recycling. Importantly, TDM1 shows similar internalization rates at late time points in control versus statin-incubated cells. This temporal *in vitro* effect of statins suggests that internalization takes place once the transient modulation has worn out.

The above data provided the rationale for preclinical imaging studies to explore the potential role of lovastatin as a CAV1 modulator in the context of TDM1 binding to HER2^+^ GC. Mice bearing subcutaneous xenografts were orally administered the previously reported dose schedule of the cholesterol-depleting drug (two doses of 8.3 mg/kg given 12 h apart) [22]. Lovastatin induced a significant reduction in CAV1 tumor levels and increased HER2 membrane levels (**Fig. 3E**). At 48 h after the first dose of lovastatin, CAV1 expression and HER2 staining resembled those found at 0 h, lending further evidence for the transience and temporality of our statin regimen. To non-invasively monitor ADC uptake in statin cohorts, mice were intravenously injected with ^89^Zr-labeled TDM1 at 12 h after the first dose of lovastatin. The 12 h window for antibody injection was based on our observations of CAV1 depletion and an enhancement in membrane HER2 (**Fig. 3E**). Control cohorts included mice orally administered saline instead of statin. The saline cohort revealed a radiopharmacologic profile standard for zirconium-89 labeled antibodies (**Fig. 3F, Supplementary Fig. 7A**) with gradual antibody accumulation to xenografts (4.9 ± 2.7, 10.2 ± 2.9, 22.2 ± 12.6, 31.1 ± 9.3 %ID/g at 4, 8, 24, and 48 h). However, the two doses of lovastatin yielded an antibody uptake higher at the different time-points when compared with the saline cohort (15.5 ± 9.7, 19.5 ± 9.3, 52.1 ± 7.6, 63.4 ± 16.7 %ID/g at 4, 8, 24, and 48 h). Oral administration of lovastatin results in images with high contrast and enhances tumor-to-background ratios (**Fig. 3F, Supplementary Fig. 7B**). To determine whether statin-mediated enhancement in ADC-tumor binding is dependent on CAV1 tumoral levels, we performed biodistribution studies with ^89^Zr-labeled TDM1 in the HER2-positive gastric PDXs shown in **Fig. 2B**. Although TDM1 accumulation in CAV1-low PDXs was similar in control and lovastatin cohorts (**Fig. 3G**), TDM1 uptake in PDX #1 (CAV1, IHC 3+) and PDX #2 (CAV1, IHC 2+) was 1.8-fold and 1.4-fold higher in lovastatin cohorts when compared with saline. These results indicate that acute CAV1 depletion by lovastatin increases cell surface receptors, enhancing TDM1 binding to xenografts with incomplete density of HER2 cell surface receptors.

**Figure 3.**
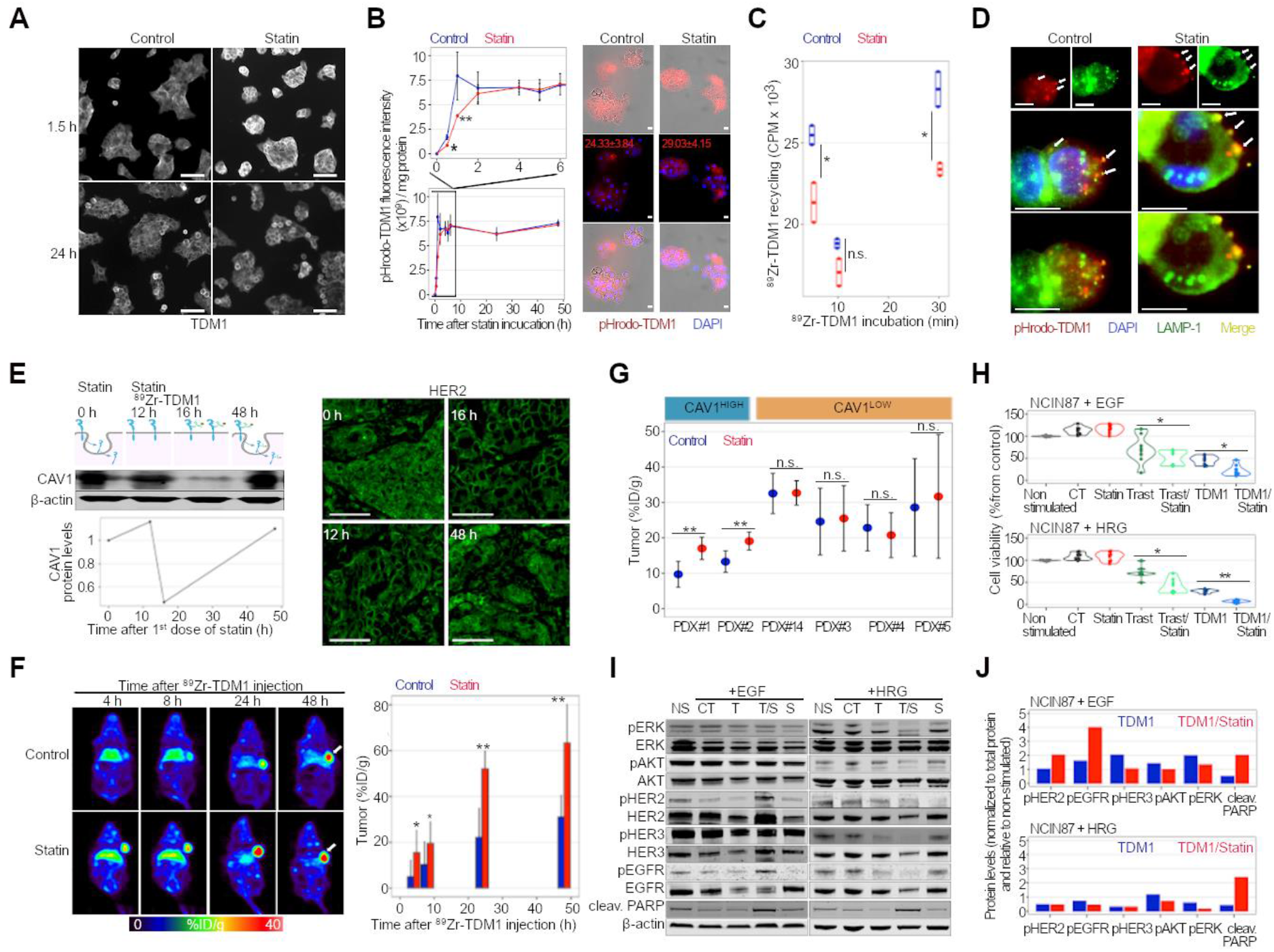
CAV1 knockdown or temporal depletion using statin enhance TDM1 binding to HER2+ CAV1-high GC. (**A**) Confocal images of immunofluorescence of TDM1 in NCIN87 cells incubated with 1 μM trastuzumab for 90 min or 24 h in the presence or absence of lovastatin. Scale bars: 20 μm. (**B**) TDM1 internalization in NCIN87 cells in the presence and absence of lovastatin. NCIN87 cancer cells were incubated with 1 μg/mL of pHrodoTDM1 for 30 min at 4°C. Cells were then released at 37°C, and the fluorescent signal was quantified at between 0.5 to 48 h after incubation with pHrodo-TDM1. The pHrodo-TDM1 fluorescent signal was normalized to the amount of protein at each time point (\**P < 0.05* and *\*\*P < 0.01* based on a Student’s *t*-test, *n* = 4). Confocal images of NCIN87 gastric cancer cells incubated with TDM1 conjugated to a red fluorescent pH-sensitive dye (pHrodo-TDM1, 1 μg/mL) for 48 h in the absence (control) or presence of lovastatin (25 μM). Scale bar: 20 μm. (**C**) TDM1 recycling in NCIN87 cells in the presence and absence of lovastatin. Cells were incubated with ^89^Zr-labeled TDM1 (1 μCi, 0.25 μg) for 4 h at 37°C. After 4 h incubation time, unbound and membrane-bound ^89^Zr-labeled TDM1 was removed, and TDM1 recycling to the cell membrane collected at 5, 10, 30 min (bars, n = 3, mean ± S.E.M, \**P < 0.05* based on a Student’s *t*-test). (**D**) Confocal images of immunofluorescence staining of pHrodo-TDM1 and LAMP1 in NCIN87 cells in the presence and absence of lovastatin. Scale bars: 100 μm and 50 μm (inset). (**E**) Western blot of CAV1 and HER2 immunofluorescence in NCIN87 s.c. tumors from athymic nude mice. Lovastatin (8.3 mg/kg of mice) was orally administrated twice with an interval of 12 h between each administration. Tumors were collected between 0 and 48 h after the first dose of lovastatin, CAV1 and HER2 were analyzed by western blot and immunofluorescence. Scale bars: 50 μm. (**F**) Representative coronal PET images and TDM1-tumor uptake at 4, 8, 24, and 48 h p.i. of [^89^Zr]Zr-DFO-TDM1 in athymic nude mice bearing s.c. NCIN87 tumors. Lovastatin (8.3 mg/kg of mice) was orally administrated 12 h prior and at the same time as the tail vein injection of [^89^Zr]Zr-DFO-TDM1 (6.66–7.4 Mbq, 45–50 μg protein). Bars, n = 5 mice per group, mean ± S.E.M. \**P < 0.05*, *\*\*P < 0.01, ***P < 0.001* based on a Student’s *t*-test. %ID/g, percentage of injected dose per gram. (**G**) [^89^Zr]Zr-DFO-TDM1 uptake in HER2-expressing gastric PDXs containing varying levels of CAV1 and administered PBS or saline. NSG mice bearing subcutaneous PDXs were intravenously administered with [^89^Zr]Zr-DFO-TDM1 (6.66–7.4 Mbq, 45–50 μg protein) and biodistribution performed at 48 h p.i. of ^89^Zr-labeled antibody. Lovastatin (8.3 mg/kg of mice) was orally administrated 12 h prior and at the same time as the tail vein injection of [^89^Zr]Zr-DFO-TDM1. PDX IDs in this figure match patient IDs shown in **Figure 1**. Points, *n* = 5 mice per group, mean ± S.E.M, *\*\*P < 0.01* based on a Student’s *t*-test. %ID/g, percentage of injected dose per gram. (**H**) Cell viability of NCIN87 cells at 48 h after cells incubation with trastuzumab and TDM1 alone or in combination with lovastatin. Cancer cells stimulated with 100 ng/mL EGF or HRG were incubated with 20 nM trastuzumab or TDM1. Lovastatin was added at a concentration of 25 μM. Non-stimulated cells were incubated in media in the absence of antibody or growth factors. Bars, *n* ≥ 3 per group, mean ± S.E.M. (**I,J**) Western blots of HER2 signaling and quantification of NCIN87 cells after 48 h incubation with TDM1 alone or in combination with lovastatin. Bars, *n* ≥ 3 per group, mean ± S.E.M.

### Statins enhance TDM1 efficacy

To assess TDM1 efficacy in combination with lovastatin, we first conducted therapy studies in CAV1-expressing HER2-positive (**Fig. 3H-J**) and HER2-negative GC cells (**Supplementary Fig. 8A**). In HER2-positive NCIN87 cells, TDM1 decreased viability (**Fig. 3H**), and lovastatin alone did not induce cell toxicity. However, statins greatly reduced cell viability when combined with the ADC. Of note, this effect was not observed in HER2-negative GC models (**Supplementary Fig. 8B,C**). Additionally, cytotoxicity was significantly higher in TDM1/statin-treated cells than cells treated with trastuzumab/statin. The increase in PARP cleavage further validated efficacy results with the combination therapy (**Fig. 3I,J; Supplementary Fig. 8D,E**). We next evaluated whether differences observed in cytotoxicity would interfere with HER2-mediated oncogenic signaling pathways. The phosphorylated proteins p-EGFR, p-ERK, and p-AKT were detected in both unstimulated and EGF-stimulated NCIN87 cells, suggesting that both MAPK and PI3K/AKT pathways are active (**Fig. 3I,J; Supplementary Fig. 8D**). TDM1 treatment alone in EGF-stimulated cells did not alter p-ERK, p-AKT, p-HER2, or p-HER3, in agreement with previous observations [42, 43]. Lovastatin did not induce significant alterations in signaling, but when combined with TDM1 it reduces phosphorylation of both ERK and AKT in EGF-stimulated cells. Under heregulin (HRG) stimulation, the ADC decreases p-ERK and p-HER and, in combination with a statin, it effectively reduces p-ERK, p-AKT, and p-HER (**Fig. 3I,J; Supplementary Fig. 8E**). Collectively, these results show that statins enhance *in vitro* TDM1 efficacy by decreasing p-ERK and p-AKT oncogenic signaling pathways.

Encouraged by the *in vitro* cell death and signaling findings, we next determined TDM1 efficacy when combined with lovastatin using NCIN87 xenografts or PDXs shown in **Fig. 1E**. Mice received intravenous injections of TDM1 (5 mg/kg once a week [42] for 5 weeks), oral doses of lovastatin (4.15 mg/kg administered 12 h prior and at the same time as the intravenous injection of antibody [22]), or a combination of ADC and lovastatin over 5 weeks (**Fig. 4A**). The vehicle and lovastatin cohorts had a similar trend of increased tumor volume over time (**Fig. 4B**). TDM1 alone inhibited tumor growth, but tumors developed resistance after 42 days of therapy. The combination of the ADC with lovastatin greatly decreased tumor volume when compared with the monotherapy. Additionally, TDM1/lovastatin decreases oncogenic signaling as we observed a reduction in p-AKT, p-ERK, and p-Tyr (**Fig. 4C, Supplementary Fig. 8F**). The phosphorylation of the cyclic (c)AMP responsive element binding protein (CREB), a player in HER2-mediated cancer development [44], was also lower in xenografts of mice treated with TDM1/lovastatin when compared with TDM1 alone (**Fig. 4C, Supplementary Fig. 8F**).

**Figure 4.**
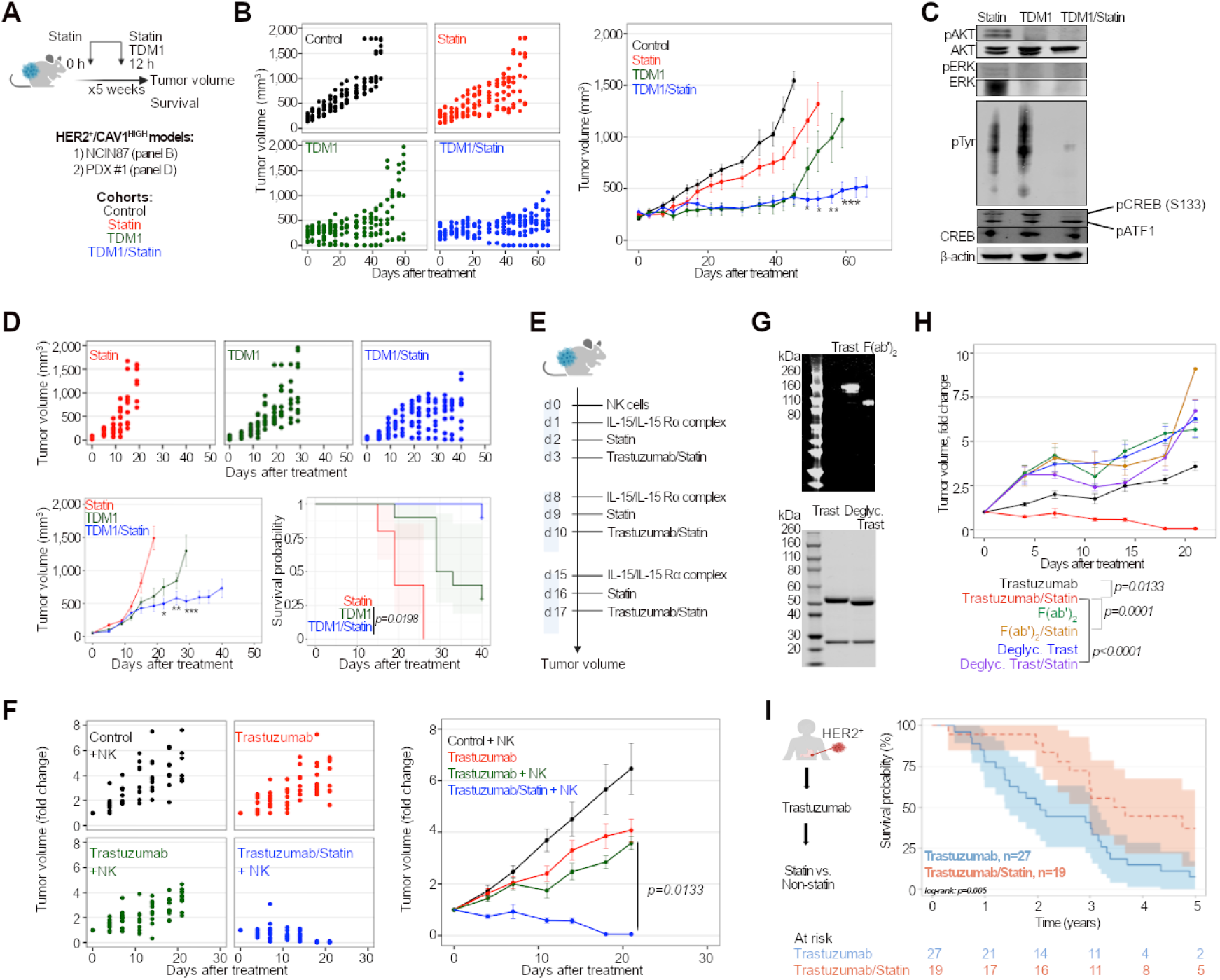
Lovastatin enhances TDM1 efficacy and trastuzumab-mediated ADCC. (**A-D**) Superior *in vivo* therapeutic efficacy of TDM1 combined with lovastatin when compared with TDM1 alone. (**A**) Intravenous TDM1 administration 5 mg/kg weekly (for 5 weeks) was started at day 0. Lovastatin (4.15 mg/kg of mice) was orally administrated 12 h prior to and simultaneously with the intravenous injection of TDM1. Lovastatin enhanced TDM1 efficacy and mice survival of *nu/nu* female mice bearing NCIN87 gastric xenografts (**B**), and NSG mice bearing CAV1-high PDXs (**D**). \**P < 0.05*, *\*\*P < 0.01, ***P < 0.001* based on a Student’s *t*-test (n ≥ 8 mice per group). (**C**) Western blot analyses of AKT, ERK, Tyr, and CREB protein expression and phosphorylation in NCIN87 xenografts at 60 days after treatment with lovastatin, TDM1, or TDM1/lovastatin. (**E,F**) NSG mice bearing NCIN87 xenografts were intravenously injected 1 × 10^6^ human NK cells at day 0. One day after NK cells tail vein injection, the IL-15/IL-15Rα complex was intraperitoneally administered at a dose of 1.25 μg/mouse. Trastuzumab or trastuzumab/lovastatin efficacy was then evaluated during a cytokine-dependent NK expansion phase (week 1 to week 3). Lovastatin enhanced trastuzumab efficacy in NSG mice humanized with NK cells and bearing NCIN87 xenografts (n ≥ 8 mice per group). (**G**) SDS-PAGE of trastuzumab, trastuzumab F(ab’)2 fragments, and deglycosylated trastuzumab. (**H**) Trastuzumab/lovastatin efficacy is higher than the combination of Fc-silent trastuzumab (trastuzumab F(ab’)2 fragments or deglycosylated trastuzumab) in NSG mice humanized with NK cells and bearing NCIN87 xenografts (n ≥ 8 mice per group). (**I**) Kaplan-Meier analysis of statin use and HER2-expressing GC disease outcome in patients treated with trastuzumab. Patients without statin treatment (blue color, *n* = 27) have a worse survival than patients treated with statin (red color, *n* = 19). *Log rank; p=0.005*.

In addition to the conventional xenografts, PDX #1 was used to validate therapeutic studies. PDX #1 was obtained from HER2^+^ GC of a patient prior initiating trastuzumab therapy. Medical records indicated that this patient underwent first-line therapy with trastuzumab but passed away less than a year after diagnosis. The patient developed brain metastases characterized by persistent *TP53* and *KRAS* somatic mutations, HER2 IHC 3+ and CAV1 IHC 3+. Lovastatin enhanced TDM1 efficacy in PDX #1, improved mice survival, and decreased p-ERK/p-AKT compared with monotherapy cohorts (**Fig. 4D, Supplementary Fig. 9A**). Notably, PDX #1 mice survival in TDM1/lovastatin cohorts was higher than trastuzumab/lovastatin (**Fig. 4D, Supplementary Fig. 9B**). These preclinical results indicate that 2-weekly doses of statin (4.15 mg/kg) given over 5 weeks to mice with CAV1-high HER2^+^ gastric xenografts enhance TDM1 efficacy.

### Statin enhances anti-HER2 antibody ADCC

Receptor internalization affects ADC efficacy (**Figs. 2–4**) and diminishes antitumor immunity by ADCC [18], a major mechanism of clinical efficacy of IgG1 therapeutic antibodies. Although antibody/lovastatin delays tumor growth in immunodeficient mice via signaling inhibition, xenograft regrowth arises in immunodeficient hosts (**Fig. 4B,D**). Because trastuzumab-mediated ADCC happens mainly through NK cells [7–9], we isolated NK cells from a healthy donor (**Supplementary Fig. 10A-D**) to measure ADCC in HER2+ GC cells expressing different levels of CAV1 (**Supplementary Fig. 4A,B**). The cholesterol-depleting drug enhanced antibody ADCC in CAV1-positive GC cells but not in CAV1-negative cells (**Supplementary Fig. 10E**). Next, we conducted therapeutic studies in NK-humanized NSG mice bearing NCIN87 xenografts (Fig. 4E). The NK cohorts showed initial response to trastuzumab, but tumor progression occurred on day 10 (**Fig. 4F, Supplementary Fig. 10F**). On the other hand, trastuzumab/lovastatin combination therapy yielded tumor regression and stabilization. Control experiments used F(ab′)2 fragments and deglycosylated trastuzumab (**Fig. 4G**) to remove the Fc-mediated stimulation of NK cell effector function. The combination of lovastatin with Fc-silent trastuzumab shows lower efficacy than trastuzumab/lovastatin (**Fig. 4H**).

To further confirm ADCC findings, we monitored trastuzumab/lovastatin efficacy during cytokine-dependent (between week 1 and week 3) and cytokine withdrawal phases (weeks 4 to 5, **Supplementary Fig. 11A**). NK cells were present on day 7 after adoptive transfer, and NK expansion occurred during the cytokine-dependent phase (**Supplementary Fig. 11B**). The number of NK cells then decreased between days 21 and 30. All mice in the trastuzumab/lovastatin cohort showed tumor regression and stabilization in tumor volume during the first three weeks (**Supplementary Fig. 11C**). In contrast, 7 out of 10 mice demonstrated tumor regrowth in the cytokine withdrawal phase. These results indicate that lovastatin enhances trastuzumab efficacy, which depends on cytokine-mediated NK expansion and antibody’s Fc domain.

### Patient survival is increased among statin users in trastuzumab GC clinical trials

Retrospective analyses of patients with gastric cancer undergoing trastuzumab therapy further validated our preclinical findings (**Fig. 4I**). This study compared investigator-assessed responses between patients with and without background statin use for overall survival. Forty-one percent of patients (19 of 46) received statin as a background prescription while receiving trastuzumab (**Table 1**). Twenty-six percent of patients were taking statins before initiating HER2-targeted treatments. Statin users were more frequently male (86% versus 14%) and more often older than 60 years of age (80% versus 20%). In the retrospective analyses, statin users treated with trastuzumab had longer survival than non-statin users (log-rank, *p=0.005;* Fig. 4I). Statin users in the CAV1-high expression group demonstrated longer survival when compared with non-statin users (log-rank, *p=0.02;* **Supplementary Fig. 12A**).

## DISCUSSION

The retrospective clinical analyses and preclinical activity of trastuzumab and TDM1, shown here, demonstrate that efficacy depends on density of the surface-localized receptors rather than the total quantity of the protein. These findings are significant in gastric tumors with heterogeneous and incomplete patterns of HER2 membrane reactivity [2, 10–12]. The variability in GC response to anti-HER2 antibody therapies among patients [5, 6], all with seemingly HER2+ disease, suggests that patient selection should be optimized by incorporating other aspects of the biology. Tumors with high levels of tumor heterogeneity in HER2 expression have a poor response to TDM1 [45–47]. CAV1 may be a complementary biomarker to detect HER2 expression in the clinic and predictive biomarker of targeted therapeutic response as determined in our retrospective screening. This led to the hypothesis that acute CAV1 depletion is a potential pharmacologic strategy to enhance HER2 at the cell membrane, contributing to a more homogeneous receptor staining and further improving response to antibody therapy. From a mechanistic perspective, acute CAV1 depletion delays ADC recycling while enhancing disruption of downstream signaling and Fc-mediated stimulation of NK cell function.

CAV1 role as a tumor suppressor or promoter depends on the tumor type and disease stage [48]. In HER2-positive tumors, recombinant overexpression of CAV1 blocks oncogenic signaling [49]. Here, we show distinctive signaling profiles in HER2^+^ CAV1-low versus CAV1-high gastric tumors. *PIK3CA* mutations, *EGFR*-and *MET*-amplification, previously demonstrated to determine clinical response to HER2-targeted therapies [15], were observed in CAV1-low GC. The low levels of p-HER2, p-ERK, and p-Tyr in CAV1-high GC support previous reports of a negative regulation between signal transduction and CAV1 protein and suggest CAV1 as a tumor suppressor [23, 49, 50]. On the other hand, low tumoral levels of CAV1 correspond to tumors with high HER2 density at the membrane of GC and might be preferable when selecting patients for anti-HER2 antibody therapies. CAV1 modulates membrane levels of HER2 within the context of receptor endocytosis [22, 26, 27], interfering with the uptake and efficacy of antibodies or ADCs [22, 24, 25, 28, 29]. This study shows that the expression of CAV1 is an independent predictor of poor overall survival during trastuzumab therapy in HER2^+^ GC.

Although TDM1 mechanisms of action are complex, the canonical model of ADC provides an helpful framework [51]: (i) ADC binds to the cell-surface receptor, (ii) ADC retains the antibody component and can decrease downstream signaling or induce ADCC, (iii) ADC-HER2 internalizes from the membrane into the intracellular compartment, and (iv) linker breakdown and DM1 drug release. Notably, the current status of patient selection for antibody-targeted therapies does not account for HER2 cellular distribution. It therefore may overestimate the amount of antigen available at the cell membrane for engaging the ADC. In this context, a lack of correlation between HER2 density and trastuzumab accumulation in tumors is reported in clinical immunoPET imaging studies [45, 52]. Others have shown that HER2 density at the cell membrane is a strong predictor of clinical outcome in patients with advanced breast cancer treated with trastuzumab and chemotherapy [53]. Our immunoPET studies using PDX models that resemble the genetic complexity and heterogeneity of GC [54] indicate low TDM1 uptake in tumors containing high CAV1 protein levels. Along similar lines, the HER2-to-CAV1 ratio is associated with GC cells’ efficiency in binding and internalizing TDM1. The impact of caveolae-mediated endocytosis has been reported for TDM1 and other ADCs: ADCs co-localize with CAV1 in resistant cancer cells [25, 55]. However, it is important to keep in mind that a decreased receptor expression preventing TDM1 from binding to GC is just one of many factors influencing antibody uptake. Other examples include truncated HER2 isoforms [56] or dysregulated mechanisms of ADC recycling, endocytosis, catabolism, and efflux of payload [17, 19, 25].

Kinase inhibitors [19, 57, 58] and CAV1 modulators [24] can temporally enhance HER2 membrane levels or promote ADC endocytosis in ways that improve TDM1 efficacy. While some reports suggest a positive role for CAV1 in promoting cells’ sensitivity to TDM1 [28, 29], others have shown an association between resistance and caveolae-mediated ADC internalization [25]. CAV1 knockdown augments cell-surface HER2 half-life [22]; here, we confirm that CAV1 downregulation improves TDM1 binding to xenografts. In contrast, increased TDM1 efficacy occurs *in vitro* after enhancing CAV1 expression by metformin [24]. Although metformin is not a specific caveolae modulator and may exert other pleiotropic biologic effects, the proposed mechanism of action in that study is that caveolae mediate TDM1 endocytosis. Considering this previous report, it would seem disadvantageous to reduce CAV1 in tumors treated with TDM1 as it could decrease the ADC internalization. However, our data show that CAV1 modulation is unlikely to change HER2 degradation processes and ADC cellular trafficking for later time points of administering a cholesterol-depleting drug. Instead, transient and controlled CAV1 depletion boosts TDM1-tumor binding while reducing ADC recycling. This is consistent with previous preclinical studies temporally increasing cell-surface receptors to potentiate TDM1 therapy [58].

Controlled use of lovastatin, at doses lower than the maximum human dose, produce a temporal decrease in CAV1 protein levels [35]. Compared with lipophilic lovastatin, hydrophilic statins are less able to cross cancer cell membranes as they require active transport to enter cells [59]. Importantly, the lipophilic prodrug lovastatin depletes tumoral CAV1 in ways that improve antibody binding to tumors [22, 34, 35]. Similarly, lovastatin enhances TDM1 uptake in CAV1-high PDXs, confirming an association between the loss of CAV1 and ADC uptake. Additionally, statins do not alter TDM1 accumulation in tumors with low levels of CAV1. Still, a large number of other proteins involved in non-caveolae internalization pathways are temporally affected by statins [35]. Nonetheless, specific genetic depletion of CAV1 *in vivo* enhances ADC accumulation in tumors. Thus, these data show that acute depletion of CAV1, whether induced by synthetic oligonucleotides or small molecules, is an important event for anti-HER2 antibody binding to GC.

After reaching tumor cells and engaging with membrane HER2, TDM1 retains the functionality of its naked antibody, trastuzumab. Therefore, the initial therapeutic mechanisms of TDM1 start before ADC internalization and payload release inside the cancer cells. From a mechanistic perspective, lovastatin enhances the Fab-mediated activity of trastuzumab responsible for disrupting oncogenic signaling. When compared with trastuzumab, statins induce higher ADC therapy, suggesting they ultimately increase DM1-mediated cytotoxicity. However, temporally enhancing cell-surface HER2 overall may not augment therapeutic outcomes if other resistance mechanisms (e.g. alterations in oncogenic signaling and tumors’ vulnerability to microtubule-directed chemotherapy) are at play [51]. In addition to the Fab region, the Fc portion of the antibody increases cell death by orchestrating ADCC. We show that CAV1 depletion boosts NK-mediated ADCC, an important mechanism of the clinical effectiveness of trastuzumab [7–9]. Others have also shown that endocytosis modulation enhances Fc-mediated ADCC of antibodies against HER or PD-L1 [18]. The increased ADCC in combination strategies using statin depends on HER2 surface levels, the Fc-domain of the antibody, and the presence of cytokine-expanded NK cells.

*In vitro* lovastatin doses reported here are higher than nanomolar range doses detected in patient prescriptions. Therefore, *in vitro* results herein described might not translate to clinical dosages. In preclinical models, lovastatin was administered on a two-dose schedule. Therefore, low and variable amounts of the cholesterol-depleting drug will accumulate and reduce tumoral CAV1. However, our retrospective findings of statin use during patient treatment with trastuzumab uphold the translational relevance of results obtained in PDXs. Note that these analyzes collected data from patients receiving standard statin doses for cardiovascular indications, while our therapeutic studies relate to mice with normal cholesterol levels. Additionally, the patient survival analyses do not account for variables that possibly cause residual confounding observations (e.g. physical activity, obesity, and diet). Further, the ability of statins to increase trastuzumab efficacy may depend on the tumor’s genotype, and future analyses with increased sample size will stratify statin users into CAV1-high versus CAV1-low tumors. Although a prospective trial is needed to confirm this combination approach fully, our preclinical and retrospective studies show statins’ potential to enhance antibody-directed therapies in GC.

An extensive pharmacodynamic/pharmacokinetic study is required for a clinical investigation of statin use in combination with HER2-targeted therapies. The outcome of these studies will grant insights into possible coincidental consequences on non-tumor cells, unsought toxicities, and statin doses for clinical use. Based on the medical records, statin users included here initiated statins before or while on trastuzumab therapy and presumably tolerated these cholesterol-depleting agents well. Other studies also suggest that statins prevent heart failure in patients receiving trastuzumab [60]. Therefore, these results indicate that the combination approach may hold a clinically acceptable safety profile and may achieve reasonable tumors selectivity when used in a controlled manner.

In summary, CAV1 may serve as a predictive biomarker when selecting tumors for HER2-targeted therapies. Importantly, immunoPET is an incredible methodology to measure differences in trastuzumab or ADC binding to CAV1-high versus CAV1-low tumors. Statin-mediated temporal increase in HER2 receptors at the cell surface has the capability to enhance trastuzumab and TDM1 efficacy. Our findings can potentially be extended to other antibody therapies and tumor types characterized by heterogeneous patterns of receptors at the cell membrane of tumor cells. These studies may help guide future clinical trials into integrating statins — forthwith available, well-tolerated, and affordable agents — for combination approaches in cancer treatment.

## METHODS

### Ethical compliance

The studies reported here were performed in compliance with all relevant ethical regulations.

### HER2 positivity, MSK-IMPACT assay, and retrospective data

HER2+ (HER2 IHC 3+, HER2 IHC 2+ and HER2:CEP17 FISH ratio ≥ 2.0 or *ERBB2*-amplified by next-generation sequencing) gastric tumor tissues were obtained from trastuzumab trials at MSK led by Y.J. (NCT02954536, IRB# 06-103, NCT01913639, NCT01522768) [15]. Patient clinical information was collected manually from the electronic medical record (M.M. and M.L.). The presence of somatic alterations in HER2-expressing tumors was analyzed by MSK-IMPACT, as previously described [15, 61].

### CAV1 immunohistochemistry (IHC) optimization and scoring

CAV1 IHC optimization was performed by the Laboratory of Comparative Pathology at MSK. Anti-CAV1 antibodies obtained from two different commercial sources (Abcam, ab2910 and BD Biosciences, 610407) were used for IHC. After comparing CAV1 IHC in various human tissues, the BD Biosciences antibody demonstrated lower unspecific reactivity than the Abcam antibody. IHC staining for CAV1 was performed on formalin-fixed paraffin-embedded HER2-expressing gastric tumor sections (5 μm thickness) on a Leica Bond RX automated stainer (Leica Biosystems, Buffalo Grove, IL). Following deparaffinization and heat-induced epitope retrieval in a citrate buffer at pH 6.0, the primary antibody against CAV1 (BD Bioscience, 610407) was applied at a concentration of 1:250 (v/v). A polymer detection system that includes an anti-mouse secondary antibody (DS9800, Novocastra Bond Polymer Refine Detection, Leica Biosystems) was then applied in the tumor samples. The 3,3’-diaminobenzidine tetrachloride (DAB) was used as the chromogen, and the sections were counterstained with hematoxylin.

Initial titration studies of CAV1 IHC optimization used human tonsil tissues (**Supplementary Fig. 1**). After CAV1 immunohistochemical validation, both anti-CAV1 Abcam ab2910 and BD Biosciences 610407 were used to stain a HER2^+^/CAV1^LOW^ and HER2^+^/CAV1^HIGH^ tumor expresser (**Supplementary Fig. 1**). The high and low CAV1 tumor expressers were based on previous IF analyses (**Supplementary Fig. 2**). The anti-CAV1 antibody from BD Biosciences demonstrated low unspecific binding (**Supplementary Fig. 1)**. The CAV1 IHC scoring was performed by a board-certified veterinary pathologist (S.M.). The pathologist performed a blind histopathological examination without prior knowledge of CAV1 IF or HER2 membrane levels. The slides were scored according to the standard IHC scoring for HER2 in human tumors. Only CAV1 reactivity associated with the membrane of neoplastic cells was considered for scoring. Cytoplasmic CAV1 reactivity in neoplastic cells, endothelial CAV1 reactivity in stromal blood vessels (**Supplementary Fig. 1B**), or CAV1 non-specific reactivity in necrotic regions (**Supplementary Fig. 1C**) was ignored for IHC scoring. Samples with CAV1 IHC 0/1+ and CAV1 IHC 2+/3+ were classified as CAV1-low and CAV1-high, respectively.

### Immunofluorescence (IF) staining of HER2 and CAV1 in tumor tissues

The MSK Molecular Cytology Core Facility performed CAV1 and HER2 immunofluorescence staining of formalin-fixed, paraffin-embedded sections (10 μM) sections. The whole slide digital images of HER2 and CAV1 staining were obtained on Pannoramic MIDI scanner (3DHistech, Hungary) at a resolution of 0.3250 μm per pixel. Regions of interest around the cells were then drawn and exported as .tif files from these scans using Caseviewer (3DHistech, Hungary.) These images were then analyzed using ImageJ/FIJI (NIH, USA) to measure fluorescence intensity after applying a median filter and background subtraction.

### Cell lines, cell culture, and treatments with lovastatin

Human gastric cancer (GC) cell lines NCIN87, AGS, SNU5, SNU1, and KATOIII, were purchased from American Type Culture Collection (ATCC). MKN45 gastric cancer cells and embryonic kidney 293 cells (HEK 293) were gifts from the Rudin Lab and Weisser Lab at MSK. All cell lines were mycoplasma-free and cultured at 37°C in a humidified atmosphere at 5% CO2 until a maximum passage of 15. The MSK integrated genomics operation core performed cell line authentication using short tandem repeat analysis.

All cell culture media were supplemented with 100 units/mL penicillin and streptomycin. NCIN87 GC cells were maintained in Roswell Park Memorial Institute (RPMI)-1640 growth medium supplemented with 10% fetal calf serum (FCS), 2 mM L-glutamine, 10 mM hydroxyethyl piperazineethanesulfonic acid (HEPES), 1 mM sodium pyruvate, 4.5 g/L glucose and 1.5 g/L sodium bicarbonate. KATOIII cells were grown in Iscove’s Modified Dulbecco Medium (IMDM) growth medium supplemented with 20% FCS and 1.5 g/L sodium bicarbonate. MKN45 were kept in RPMI-1640 supplemented with 2 mM L-glutamine. SNU1 cells were maintained in RPMI-1640 containing 10% FCS. AGS cells were grown in Kaighn’s Modification of Ham’s F-12 Medium (F-12K Medium) supplemented with 10% FCS. SNU5 cells were grown in IMDM containing 20% FCS.

For *in vitro* experiments with lovastatin, cells were incubated with 25 μM of the active form of lovastatin (Millipore) for 4 h prior addition of trastuzumab or TDM1 [22].

### Generation of GC lines stably expressing HER2 (LV-HER2)

The pHAGE-*ERBB2* (Addgene plasmid 116734) was a gift from Gordon Mills and Kenneth Scott; lentiviral envelope and packaging plasmids pMD2.G (Addgene plasmid 12259) and psPAX2 (Addgene plasmid 12260) were gifts from Didier Trono. The plasmids were purified using QIAquick Spin Miniprep or Plasmid Plus Midi kits (Qiagen) and verified by Sanger sequencing (Genewiz) before lentiviral production. Lentivirus was produced by transfection of HEK293T cells using the JetPrime system (Polyplus). The ratio of pMD2.G:psPAX2:pHAGE-*ERBB2* was 1:2:3, the ratio of JetPrime transfection reagent to DNA was 2:1, and the ratio of JetPrime buffer:transfection reagent was 50:1. The HEK293T cells were incubated with the DNA and transfection reagents for 24 h before the media was changed. Two days after replacing the media, the media (herein referred to as viral supernatant) was collected and filtered through 0.45 μM PVDF filters (Millipore). The viral supernatant was then concentrated 20-fold with Lenti-X Concentrator (Clontech) according to the manufacturer’s instructions. The GC cell lines KATOIII, MKN45, SNU5, AGS, and SNU1 were transduced using 8 μg/mL hexadimethrine bromide (Sigma), and the media was changed 24 h after transduction. Three days after transduction, puromycin selection of HER2-expressing cells was initiated on all cell lines at concentrations from 1–2.5 μg/mL, and selection was continued for at least four days. The overall increase in HER2 cellular expression was validated by Western blot (**Supplementary Fig. 4B**).

### Preparation of human NK cells

An 81 mL leukapheresis pack containing 5.50 × 109 white cells with a viability of 98% was shipped at ambient temperature from STEMCELL Technologies (**Supplementary Fig. 10A**) and processed immediately upon receipt. On the day of arrival, the COVID-19 PCR result was pending for the donor. Therefore, samples were handled following Biosafety guidelines at MSK for human samples of unknown COVID-19 status of source case. Because the procedures were non-aerosol generating, the samples were handled as BSL2. Upon removing the supernatant, the leukapheresis sample was washed by adding 81 mL of EasySep buffer (20144, STEMCELL). The sample was then centrifuged at 500×*g* for 10 min at room temperature. Upon removal of the supernatant, the cell pellet was resuspended in EasySep buffer at 5.50 × 10^7^ cells/mL. An ELISA test for COVID-19 was performed before isolation of NK cells using the KT-1032 EDI™ Novel Coronavirus COVID-19 Elisa kit. After confirming that the sample was COVID-19 negative, the NK cells were isolated by negative selection using the human NK cell enrichment kit (19055, STEM Cell). Briefly, the enrichment cocktail (50 μL/mL) was added to the sample containing 5.50 × 10^7^ cells/mL and incubated for 10 min at room temperature. The magnetic particles (100 μL/mL) were then added and incubated for 10 min at room temperature. The sample was placed in the EasySep magnet for 10 min. The isolated NK cells were then transferred into a new tube and the NK cell population was confirmed by FACs as CD3^-^CD56^+^ cells (**Supplementary Information 10C,D**).

### Flow cytometry

After NK cell isolation, cells were washed twice with ice-cold PBS. NK cells were then split into groups and stained with APC-hCD56 (clone HCD56, 318309, Biolegend) and PE-hCD3 (clone HIT3a, 300307, Biolegend). After 20 minutes of incubation, NK cells were washed with PBS containing 2% (v/v) FBS and fixed in 4% paraformaldehyde (PFA). NK cells were then resuspended in FACS buffer (PBS containing 2% FBS and 2 mM EDTA) and placed on ice until analysis. Single color controls were made, NK cells were identified as CD3-CD56+, and results were analyzed with Flowjo software (Flowjo LLC v10.7.1). Flow cytometry was performed in the MACSQuant Analyzer 10.

### Western blots

Whole-protein extracts from cells or tumors were prepared after cell scrapping or tissue homogenization, respectively, in RIPA buffer and separated on SDS-PAGE gels (NuPAGE 4– 12% Bis-Tris Protein Gels, Invitrogen). Membranes were probed using the following primary antibodies: rabbit anti-CAV1 1:500 (Abcam, ab2910), rabbit anti-HER2 1:800 (Abcam, ab131490), mouse anti β-actin 1:20,000 (Sigma, A1978), rabbit anti-ubiquitin 1:1,000 (Cell Signaling Technology, 3933S), mouse anti-ERK 1:100 (Invitrogen, 14-9108-80), rabbit anti-pERK 1:500 (Invitrogen, 700012), rabbit anti-AKT 1:1,000 (Cell Signaling Technology, 9272S), rabbit anti-pAKT, 1:2,000 (Cell Signaling Technology, 4060S), rabbit anti-cleaved PARP, 1:1,000 (Cell Signaling Technology, 9541S), rabbit anti-pHER2, 1:500 (Abcam, ab53290), rabbit anti-HER3, 1:500 (Abcam, ab32121), rabbit anti-pHER3, 1:2,500 (Abcam, ab76469), rabbit anti-EGFR 1:1,000 (Abcam, ab52894), rabbit anti-pEGFR 1:500 (Abcam, ab40815), mouse anti-pTyr 0.5 μg/mL (EMD Millipore, 05-321X), rabbit anti-CREB 1:1,000 (Cell Signaling Technology, 9197S), rabbit anti-pCREB 1:1,000 (Cell Signaling Technology, 9198S).

The membranes were then incubated with secondary antibodies IRDye^®^800CW anti-rabbit or anti-mouse IgG 1:15,000 (LI-COR Biosciences) and imaged on the Odyssey Infrared Imaging System (LI-COR Biosciences) followed by densitometric analysis.

### PathScan antibody array kit

The MAPK Phosphorylation Antibody Array (Abcam, ab211061) was used to determine MAPK signaling changes. Total tissue lysates (500 μg) were loaded in the membranes according to the manufacturer’s instructions. The membrane arrays were then incubated with the detection antibody cocktail, and the HRP-Anti-Rabbit IgG was used to detect bound proteins. The proteins were visualized using the detection buffer mixture on a chemiluminescent blot documentation system consisting of x-ray film with a film processor followed by densitometric analysis.

### HER2 immunoprecipitation

NCIN87 cancer cells were incubated in media with 5% (v/v) of FBS in the presence of 10 μM of the proteasome inhibitor MG-132 (Sigma-Aldrich). Cells were incubated with 10 μg/mL of TDM1 in the presence and absence of lovastatin at 37°C for 4 h. Cells were then washed with cold PBS and lysed with NP-40 buffer (150 mmol/L NaCl, 10 mmol/L Tris pH 8, 1% NP-40, 10% glycerol). Forty micrograms of proteins were used as total lysates. For immunoprecipitation, protein lysates (500 μL of NP-40 buffer containing 200 μg of protein) were incubated with 10 μg of primary antibody Neu (F-11) agarose conjugate (sc-7301; Santa Cruz Biotechnology) overnight at 4°C with gentle rotation. The pellet containing the immunoprecipitated fraction was collected by centrifugation at 1,000×*g* for 30 s at 4°C, washed 3 times with NP-40 buffer and once using nuclease-free sterile water before resuspension in Laemmli buffer.

### Cell viability and HER2 signaling analyses

Cell viability was determined in cells treated with trastuzumab, TDM1, trastuzumab/lovastatin, or TDM1/lovastatin. Cells were plated in a 96-well plate (1 × 10^4^ cells/well) and pre-cultured for 24 h. Cells stimulated with 100 ng/mL of EGF or HRG Cells were incubated with 20 nM of trastuzumab or TDM1 in the absence or presence of lovastatin. Cell viability was measured at 48 h after treatments using thiazolyl blue tetrazolium bromide (MTT, Sigma). The optical density value was read at 570 nm using the Spectra Max ID5 (Molecular Devices). The percentage of cell viability was indicated by comparison with cells in the absence of stimulation or treatments. In Western blot assays of HER2 signaling, cells were plated in a 6-well plate (1 million cells/well). The day after, cells stimulated with EGF or HRG were incubated with trastuzumab, TDM1, trastuzumab/lovastatin, or TDM1/lovastatin. Total cell extracts were collected 48 h after cells’ treatment and analyzed by Western blot.

### *In vitro* therapeutic ADCC

Cells were plated at a 50:1 effector (NK):target (GC) ratio in serum-free cell culture medium supplemented with 0.1% BSA. Cells were treated with 100 μg/mL of trastuzumab or TDM1 in the absence or presence of lovastatin. After 6 h of incubation time, cell death was measured by determining LDH release using the Cytotoxicity Detection Kit (LDH; Roche).

### Tetracycline-inducible shRNA CAV1 expression (Tet-On system)

A panel of 5 different shRNA against CAV1 and 1 NTC shRNA were generated by the Gene Editing & Screening Core at MSK and cloned into the LT3GENIR4(pRRL) vector. This backbone contains a neomycin selection and an inducible Dox system (Tet-On, **Supplementary Fig. 5A**). The viral particles were produced using ExtremeGene HP (Roche) and 293FT packaging cells using a 3^rd^ generation lentivector packaging system (3 vector system). The NCIN87 cells were then infected for 24 h. After 24 h, fresh media was added to the cells. After 48 h from the infection, antibody selection was initiated at 1200 μg/ml of neomycin and kept for 2 weeks. The cells were then placed on Dox for 48 h at a concentration of 1 μg/mL to induce GFP expression and CAV1 knockdown before sorting for GFP. Dox was removed from the media, and cells were expanded for 10 days in the absence of Dox (to return CAV1 expression to baseline levels and diminish GFP expression). The overall decrease in CAV1 expression after cells incubation with Dox was validated by Western blot (**Supplementary Fig. 5B**).

### Conjugation and radiolabeling of TDM1

TDM1 was obtained from the MSK Hospital Pharmacy. The pHrodo-TDM1 was obtained by conjugating the free lysine residues of TDM1 with the amine-reactive pH-sensitive pHrodo iFL Red STP ester dye (ThermoFisher Scientific, P36014) according to the manufacturer’s instructions.

To prepare [^89^Zr]Zr-DFO-TDM1, TDM1 was first conjugated with the bifunctional chelate *p*-isothiocyanatobenzyl-desferrioxamine (DFO-Bz-NCS; Macrocyclics, Inc) and then labeled with zirconium-89 (^89^Zr), as previously described [62]. [^89^Zr]Zr-DFO-TDM1 radiochemical purity (RCP) was determined by instant thin-layer chromatography. The [^89^Zr]Zr-DFO-TDM1 conjugates used for *in vitro* and *in vivo* studies had a RCP of 99%, radiochemical yields ranging from 92–97%, specific activities in the range of 21.98–24.73 Mbq/nmol, and immunoreactivities above 90%.

### TDM1 binding, internalization, and recycling assays

For the binding assays, solutions of 89Zr-labeled TDM1 (4 μCi/μg) were prepared in PBS (pH 7.5) containing 1% w/v human serum albumin (HSA, Sigma) and 0.1% w/v sodium azide (NaN_3_, Acros Organics). Cells (1 million) were incubated with 1 μCi (0.25 μg) of the radiolabeled antibody for 1 h at 4°C. Unbound radioactivity was removed, and cells were washed twice with PBS by centrifugation. The pellet-bound radioactivity was measured on a gamma counter calibrated for zirconium-89.

For internalization assays, cells were plated in a 96-well plate (50,000 cells/well). The day after, cells were incubated with 5 nM of pHrodoTDM1 for 30 min at 4°C. Cells were then incubated at 37°C, and fluorescent measurements were performed between 30 min and 48 h hours after incubation with pHrodo-TDM1. Fluorescence was recorded using Spectra Max ID5 (Molecular Devices) at excitation wavelength 560 nm/emission wavelength 585 nm. Protein concentration was determined using the Pierce BCA Protein Assay Kit (ThermoFisher Scientific), and the pHrodo-TDM1 fluorescent signal was normalized to the amount of protein at each time point.

For recycling assays, cells were plated in a 6-well plate (1 million cells/well). The day after, cells were incubated with 1 μCi (0.25 μg) of the radiolabeled antibody in media at 37°C. After 4 h incubation time, cells were kept in ice, washed with ice-cold PBS, and the supernatant was collected. Cell surface-bound radiotracer was collected by cells incubation at 4°C for 5 min in 0.2 M glycine buffer containing 0.15 M NaCl, 4 M urea at pH 2.5. The cells were then incubated in media at 37°C to allow recycling processes. Antibody recycling to the cell membrane was measured at 5, 10, and 30 min after washing cells and collecting the cell surface-bound radiotracer. The radioactive fractions were measured for radioactivity on a gamma counter calibrated for zirconium-89.

### Immunofluorescence assays of TDM1

For immunofluorescence assays, cells were plated at 0.1 million cells/slide in chamber slides (154526, ThermoFisher Scientific) for 24 h. Cells were then incubated with 1 μM TDM1 for 90 min or 24 h at 37°C. Cells were fixed with 4% PFA, permeabilized with 1% Triton X-100 in PBS (pH 7.4) and blocked with 5% bovine serum albumin in PBS buffer, before incubation with the DAPI and secondary goat anti-human IgG fluorescently labeled with Alexa Fluor 488 (A-11013, ThermoFisher Scientific).

For immunofluorescence assays of GC cells with pHrodo-TDM1 and LAMP-1, cells grown in chamber slides were incubated with 1 μg/mL of pHrodo-TDM1 for 48 h. After cells fixation with PFA and permeabilization using 1% Triton X-100, cells were incubated with a rabbit anti-LAMP-1 primary antibody (ab24170, Abcam). Cells were then incubated with DAPI and secondary goat anti-rabbit IgG fluorescently labeled with Alexa Fluor 488 (A-11008, ThermoFisher Scientific).

### Antibody deglycosylation and F(ab′)2 fragments generation

Trastuzumab deglycosylation was achieved by adding 1.1 units of recombinant PNGaseF enzyme (New England BioLabs) per 1 μg of antibody. Trastuzumab (3 mg, 144 μL) was mixed with 3,000 units of PNGaseF enzyme (13. 3 μL from a stock solution containing 225 U/μL), 25 μL 500 mM sodium phosphate (pH 7.5), and 31.7 μL of water. The reaction was incubated at 37°C for 2 h. To remove the PNGaseF enzyme from the reaction mixture and purify deglycosylated trastuzumab, chitin magnetic beads (100 μL, E8036S, New England Biolabs) were added to the reaction mixture.

The F(ab′)_2_ fragments were generated using trastuzumab and the F(ab′)_2_ fragmentation kit following the manufacturer’s instructions (G-Biosciences).

### Tumor xenografts and patient-derived xenografts (PDXs)

The experimentation involving animals followed the guidelines approved by the Research Animal Resource Center and Institutional Animal Care and Use Committee at MSK (New York, NY), the ARRIVE guidelines, and the guidelines for the welfare and use of animals in cancer research. NCIN87, NCIN87 shRNA NTC, NCIN87 shRNA 486, or NCIN87 shRNA 479 cancer cells were subcutaneously implanted in female athymic nude mice *nu/nu* (8 to 10 weeks old, Charles River Laboratories). A total of 5 million cells were suspended in 150 μL of a 1:1 v/v mixture of medium with reconstituted basement membrane (BD Matrigel, BD Biosciences) and injected subcutaneously in each mouse.

PDX models were established by the Anti-tumor Assessment Core, from tumor specimens collected under an approved institutional review board protocol by the Research Animal Resource Center and Institutional Animal Care and Use Committee at MSK, NY [54]. Briefly, tumors were minced, mixed with Matrigel, and implanted subcutaneously in 6–8-week-old NSG mice (Jackson Laboratories).

The tumor volume (V/mm^3^) was estimated by external vernier caliper measurements, as previously described [22].

### PDX genetic and immunohistochemical validation

To confirm that PDXs herein used recapitulate parent tissue, MSK-IMPACT data were obtained in both PDX and human tumor tissues. Given that patient-derived EBV-positive lymphomas are often observed in PDX models using NSG mice [38, 39]. H&E and IHC stained sides were reviewed by a board-certified veterinary pathologist (S.M.) to exclude lymphomas in PDX models. IHC was performed by the Laboratory of Comparative Pathology at MSK for pancytokeratin, (primary antibody Dako Z0622 applied at 1:500 concentration), human CD45 (Dako M0701, 1:100), and human CD20 (Dako M0755, 1:1000) on the Leica Bond RX automated staining as previously described for the CAV1 IHC method. Carcinomas were confirmed by IHC as pancytokeratin^+^/CD45^-^/CD20^-^. B cell lymphomas excluded from preclinical studies (*n*=13) were IHC pancytokeratin^-^/CD45^+^/CD20^+^.

### CAV1 modulation using genetic and pharmacologic approaches

For preclinical imaging studies using the Tet-On system, mice were randomly assigned into the following groups (*n* = 4 mice per group): *OFF DOX*, daily oral administration of PBS for 11 days prior to tail vein injection of 89Zr-labeled TDM1; *ON DOX*, daily oral administration of 10 mg/mL of Dox for 11 days prior to tail vein injection of 89Zr-labeled TDM1; *ON/OFF DOX,* daily oral administration of 10 mg/mL of Dox for 7 days followed by oral administration of PBS for 4 days before tail vein injection of 89Zr-labeled antibody.

For preclinical imaging studies using lovastatin, mice were assigned into the following groups (*n* = 4 mice per group) [22, 35]: *Control*, oral administration of PBS 12 h prior to and at the same time as the tail vein injection of ^89^Zr-labeled TDM1; *Lovastatin*, oral administration of lovastatin (8.3 mg/kg of mice) 12 h prior to and at the same time as the tail vein injection of ^89^Zr-labeled TDM1.

### Small-animal PET and acute biodistribution studies

Mice bearing subcutaneous xenografts or PDXs (100–150 mm3 in tumor volume) were randomized before administering [89Zr]Zr-DFO-TDM1 (6.66–7.4 Mbq, 45–50 μg protein) by tail vein injection. PET imaging (*n* = 3 mice per group) and *ex vivo* biodistribution (*n* = 5 mice per group) were performed according to previously reported methods [22, 34, 35]. PET images were analyzed using ASIPro VM software (Concorde Microsystems). Radioactivity present in each organ was expressed as the percentage of injected dose per gram of organ (% ID/g).

### *In vivo* therapeutic efficacy

Mice with subcutaneous xenografts or PDXs of volume between 100 to 300 mm3 were randomly grouped into treatment cohorts (*n* ≥ 8 per group): control, TDM1, trastuzumab, lovastatin, TDM1/lovastatin, or trastuzumab/lovastatin. Mice received weekly intravenous injections of TDM1 (5 mg/kg) or intraperitoneal injections of trastuzumab (5 mg/kg) for 5 weeks. Lovastatin (4.15 mg/kg of mice) was orally administered 12 h prior and at the same time as the intravenous injection of TDM1. Tumor volumes were determined twice a week.

### *In vivo* therapeutic ADCC

NCIN87 GC cells (5 million cells suspended in 150 μL of a 1:1 v/v mixture of medium with reconstituted basement membrane) were subcutaneously implanted in female severely immunodeficient NSG (6–8 weeks old, Jackson Laboratories). Once NCIN87 GC tumor volumes reached 100 to 150 mm^3^, freshly isolated NK cells (1 million cells in 200 μL PBS) were administered by tail vein injection. The interleukin-15/ interleukin-15 receptor alpha complex (IL-15/IL-15Rα complex) was used to achieve NK cell expansion and activation *in vivo* [63, 64]. One day after NK cells tail vein injection and once per week, the IL-15/IL-15Rα complex was intraperitoneally administered at a dose of 1.25 μg/mouse. Mice were randomly grouped into treatment cohorts (*n* ≥ 8 per group): saline, lovastatin, trastuzumab, trastuzumab/lovastatin. Mice received weekly intraperitoneal injections of trastuzumab (5 mg/kg). Lovastatin (4.15 mg/kg of mice) was orally administered 12 h prior and at the same time as the intraperitoneal injection of trastuzumab. Control cohorts included treatments in NSG mice that were not intravenously administered NK cells. Additional control experiments were performed using Fc silent deglycosylated trastuzumab and trastuzumab F(ab′)2 fragments. Tumor volumes were determined twice a week.

### Quantification and statistical analyses

Data were analyzed using R v3.6.0. (http://www.rstudio.com/) or GraphPad Prism 7.00 (www.graphpad.com). Statistical differences between mean values were determined using analysis of variances (ANOVA) coupled to Scheffé’s method or a Student’s t-test. To compare treatments between cell lines, the Wilcoxon-Mann-Whitney test was performed using a 1-sided alpha of 0.05. Spearman correlation was used for correlating HER2 or HER2/CAV1 protein levels with TDM1 internalization. The overall patient survival is defined as the time from diagnosis to death. Patients alive are censored at their date of last follow-up. Survival rates are estimated using Kaplan-Meier estimator, and curves are compared using the log-rank test.

## Data availability

All data generated or analyzed during this study are included in this published article (and its supplementary information files).

## ACKNOWLEDGEMENTS

We gratefully acknowledge Dr. Ricardo D’Oliveira Albanus from Department of Computational Medicine & Bioinformatics, University of Michigan for assistance in RStudio analyses. We would also like to acknowledge Dr. Marco Russo and Daniel Zakheim from the Gene Editing & Screening Core at MSK to assist in the Tet-on system. We are grateful to Dr. Elisa De Stanchina and all the team at the Antitumor Assessment Core for helping with the PDX models. We thank Dr. Fiona Simpson and Dr. Joseph Sun insightful suggestions regarding the experiments with NK cells.

## AUTHOR’S CONTRIBUTIONS

**Conception and design:** P. M. R. Pereira, J. S. Lewis

**Development of methodology:** P. M. R. Pereira, K. Mandleywala, S. Monette, M. Cornejo, A. K. Tully, A. Ragupathi, M. Mattar, Y. Y. Janjigian

**Acquisition of data (provided animals, acquired and managed patients, provided facilities, etc.):** P. M. R. Pereira, K. Mandleywala, S. Monette, M. Lumish, K. Tully, M. Cornejo, A. Mauguen, A. Ragupathi, M. Mattar, Y. Y. Janjigian, J. S. Lewis

**Analysis and interpretation of data (e.g. statistical analysis, biostatistics, computational analysis):** P. M. R. Pereira, K. Mandleywala, S. Monette, M. Lumish, K. Tully, M. Cornejo, A. Mauguen, A. Ragupathi, M. Mattar, Y. Y. Janjigian, J. S. Lewis

**Writing, review, and/or revision of the manuscript:** P. M. R. Pereira, K. Mandleywala, S. Monette, M. Lumish, K. Tully, M. Cornejo, A. Mauguen, A. Ragupathi, M. Mattar, Y. Y. Janjigian, J. S. Lewis

**Administrative, technical, or material support (i.e. reporting or organizing data, constructing databases):** P. M. R. Pereira, K. Mandleywala, S. Monette, M. Lumish, K. Tully, M. Cornejo, A. Mauguen, A. Ragupathi, M. Mattar, Y. Y. Janjigian, J. S. Lewis

**Study supervision:** P. M. R. Pereira, Y. Y. Janjigian, J. S. Lewis

