## Supplementary Figures for "Caveolin-1 temporal modulation enhances trastuzumab and trastuzumab-drug conjugate efficacy in heterogeneous gastric cancer"

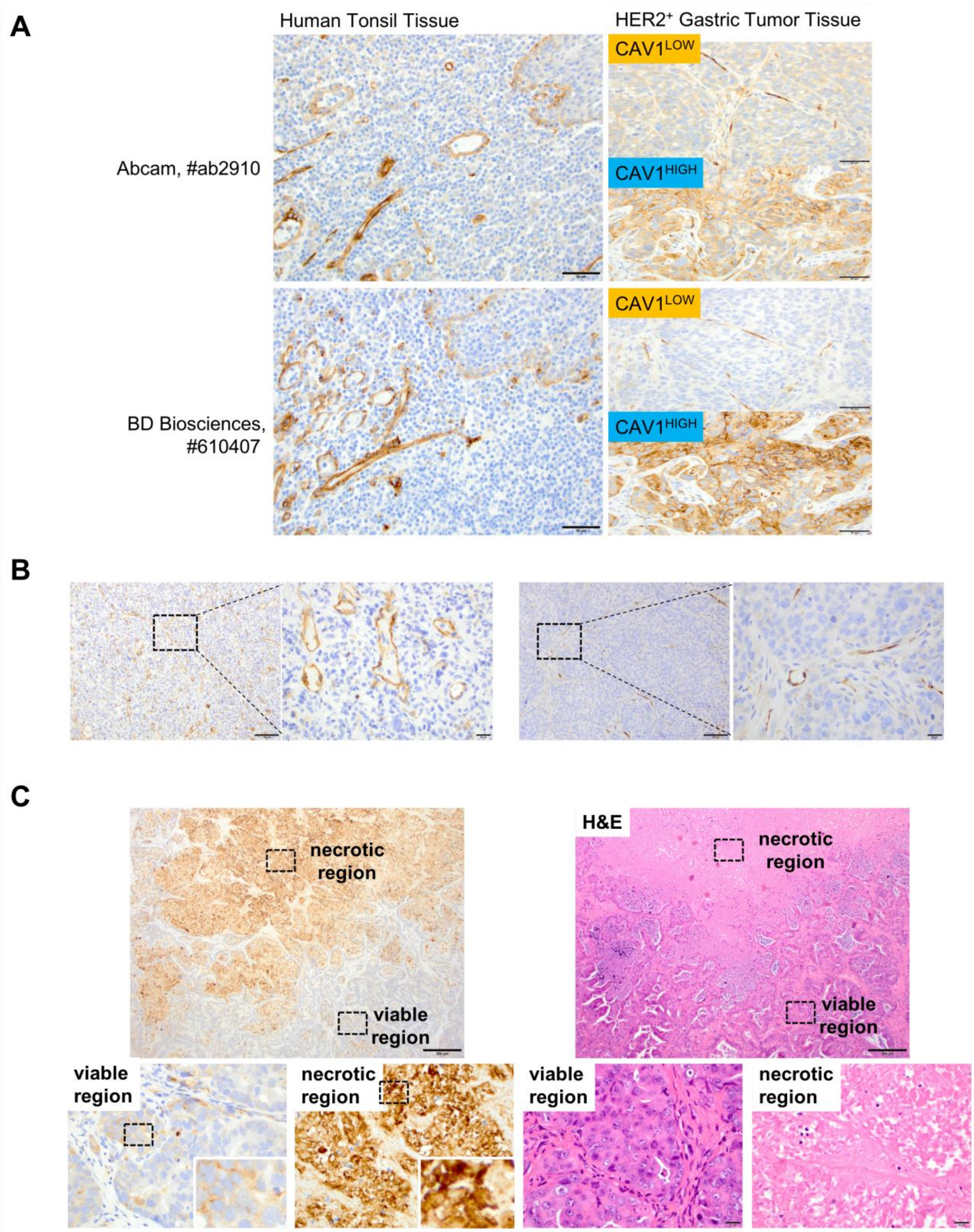


**Supplementary Figure 1.** (**A**) Immunohistochemical detection of CAV1 in human tonsil tissues (left panel) and HER2-expressing gastric tumor tissues using anti-CAV1 antibodies from Abcam (ab2910) or BD Biosciences (610407). CAV1 reactivity is present mainly in the blood vessels of human tonsils with weak epithelial reactivity. In gastric tumor tissues, CAV1 is present in both tumor and vascular endothelium. The anti-CAV1 antibody from BD Biosciences was used in future staining of HER2-expressing gastric tumor tissues as this was the one with the lowest unspecific binding. Scale bar, 50 µm. (**B**) Representative images (scale bar, 100 µm) with magnification (scale bar, 20 µm) of CAV1 staining in the vascular endothelium but not in tumor cells. CAV1 IHC scoring analyses excluded staining in the vascular endothelium. (**C**) Representative images of CAV1 IHC 1+ membrane associated staining of tumor cells in viable regions versus strong nonspecific staining of necrotic tumor areas. CAV1 staining in necrotic regions was excluded during the IHC scoring analyses. Scale bar, 200 µm.

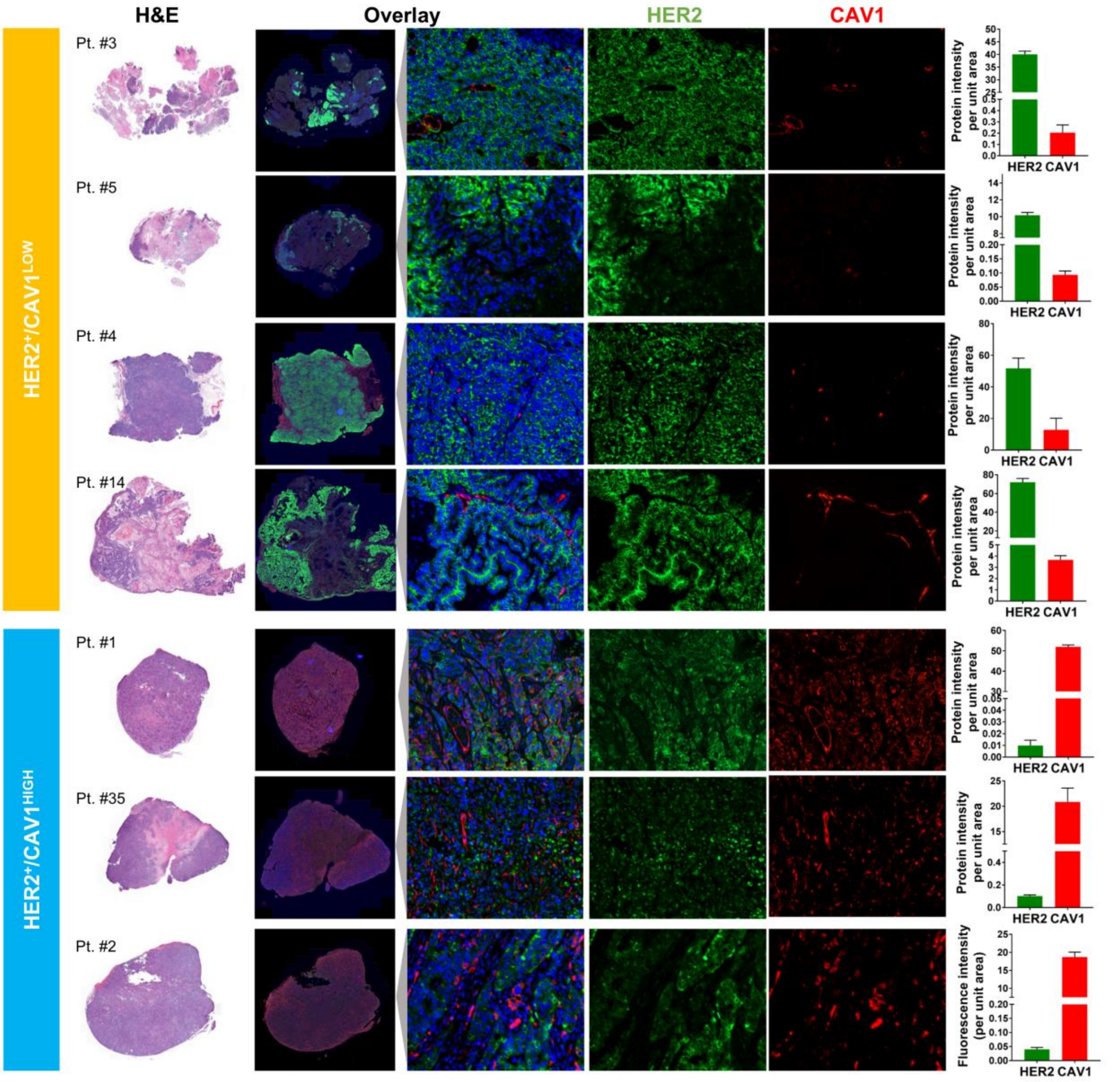


**Supplementary Figure 2.** Hematoxylin & Eosin (H&E), confocal images, and quantification of immunofluorescence staining of HER2 (green color) and CAV1 (red color) in human HER2-expressing gastric tumors. In blue color is cell nuclei. Tumors were classified as CAV1-high (Patient #1, #35, and #2) and CAV1-low (Patients #3, #5, #4, and #14). HER2 IF shows a predominant presence at the cell membrane in CAV1-low tumors. The graphs plot protein intensity per unit area, calculated by quantifying IF images (mean ± S.E.M, n = 3).

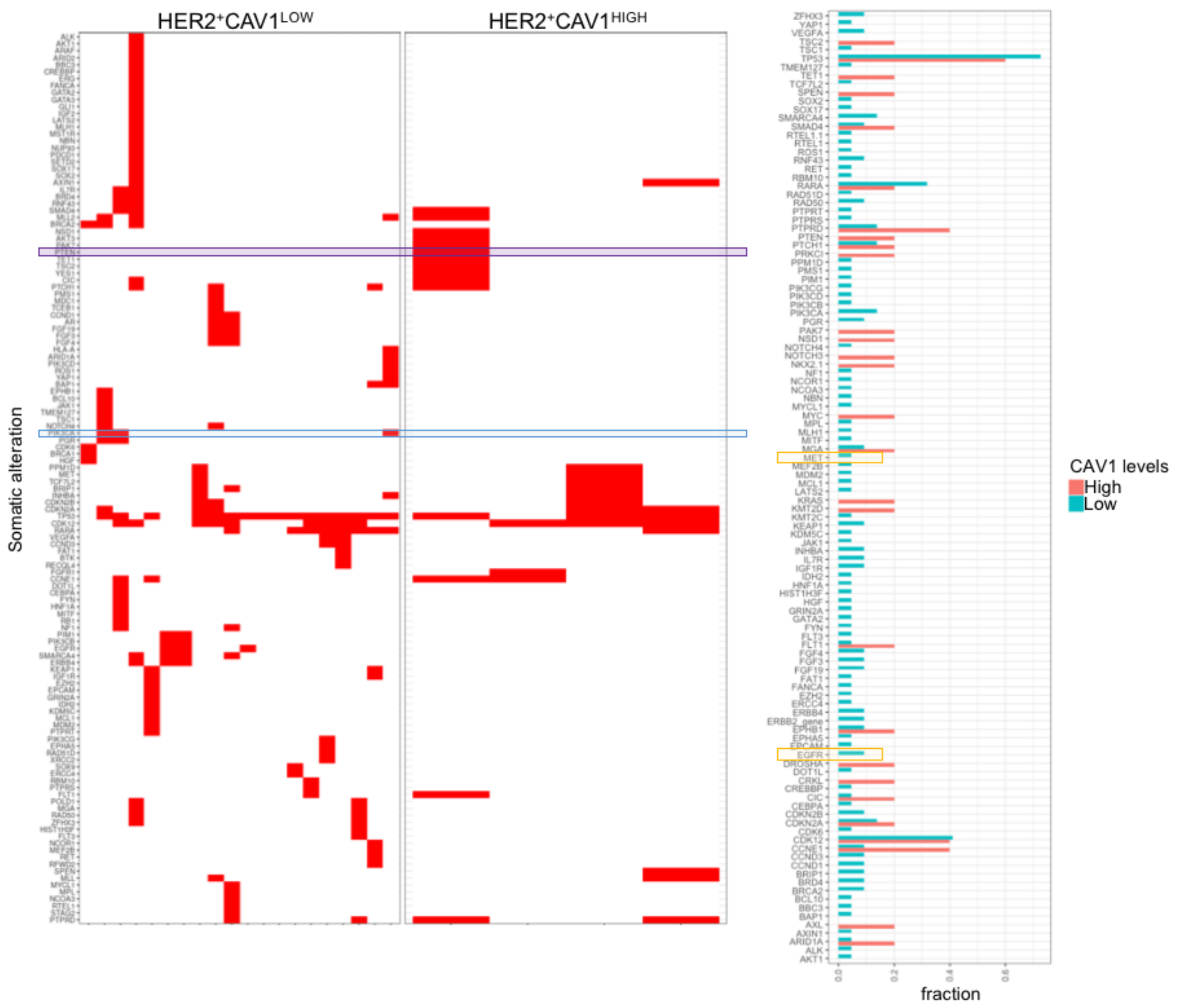


**Supplementary Figure 3.** Genomic alterations found in HER2$^+$CAV1$^{LOW}$ ($n$ = 20) versus HER2$^+$CAV1$^{HIGH}$ ($n$ = 7) gastric tumor tissues. (**Left Panel**) The rectangles with blue color highlight the absence of *PIK3CA* mutations in CAV1-high tumors. The rectangles with purple color highlight the absence of *PTEN* alterations in CAV1-low tumors. (**Right Panel**) The rectangles with yellow color highlight the presence of *MET* and *EGFR* amplification in CAV1-low tumors.

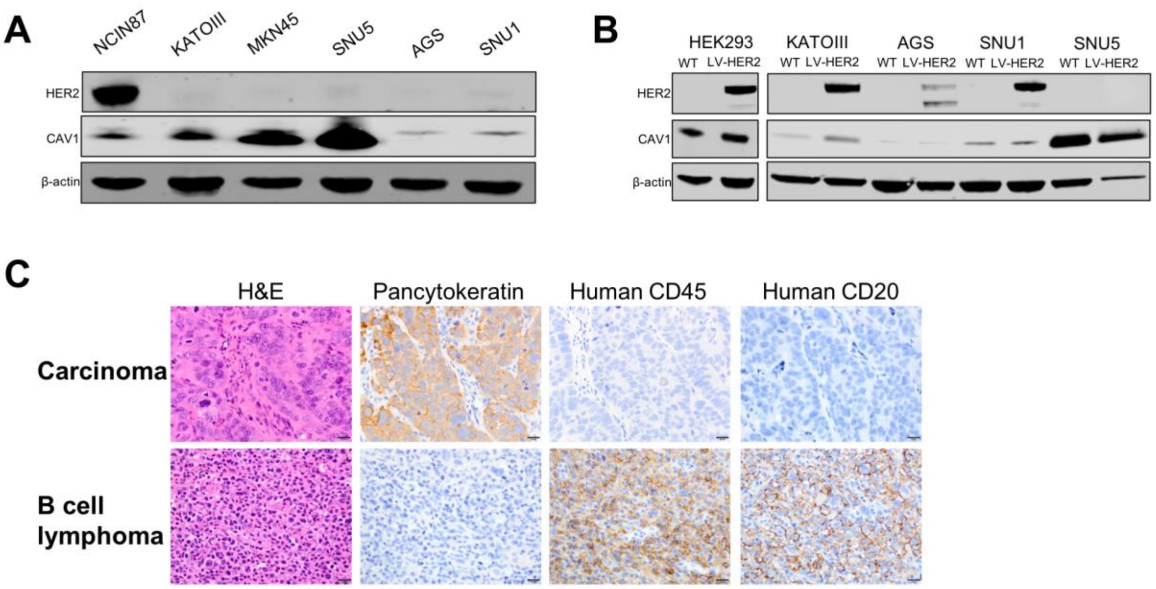


**Supplementary Figure 4.** (**A**) Western blot of HER2 and CAV1 in the total lysates of NCIN87, KATOIII, MKN45, SNU5, AGS, and SNU1 gastric cancer cells. (**B**) Western blot of HER2 and CAV1 in KATOIII, AGS, SNU1, and SNU5 gastric cancer cell lines wild-type (WT) and sublines stably expressing HER2 (LV-HER2). HEK293 WT and HEK293 LV-HER2 are also shown as controls. The protocols herein used did not allow for the successful generation of LV-HER2 in MKN45 and SNU5 cell lines containing the highest CAV1 expression. **(C)** Hematoxylin & Eosin (H&E) and immunohistochemical detection of pancytokeratin, human CD45, and human CD20 in PDX samples. Carcinomas were confirmed by IHC pancytokeratin$^+$/CD45$^-$/CD20$^-$. B cell lymphomas excluded from the studies were IHC pancytokeratin$^-$/CD45$^+$/CD20$^+$. Scale bar, 20 μm.

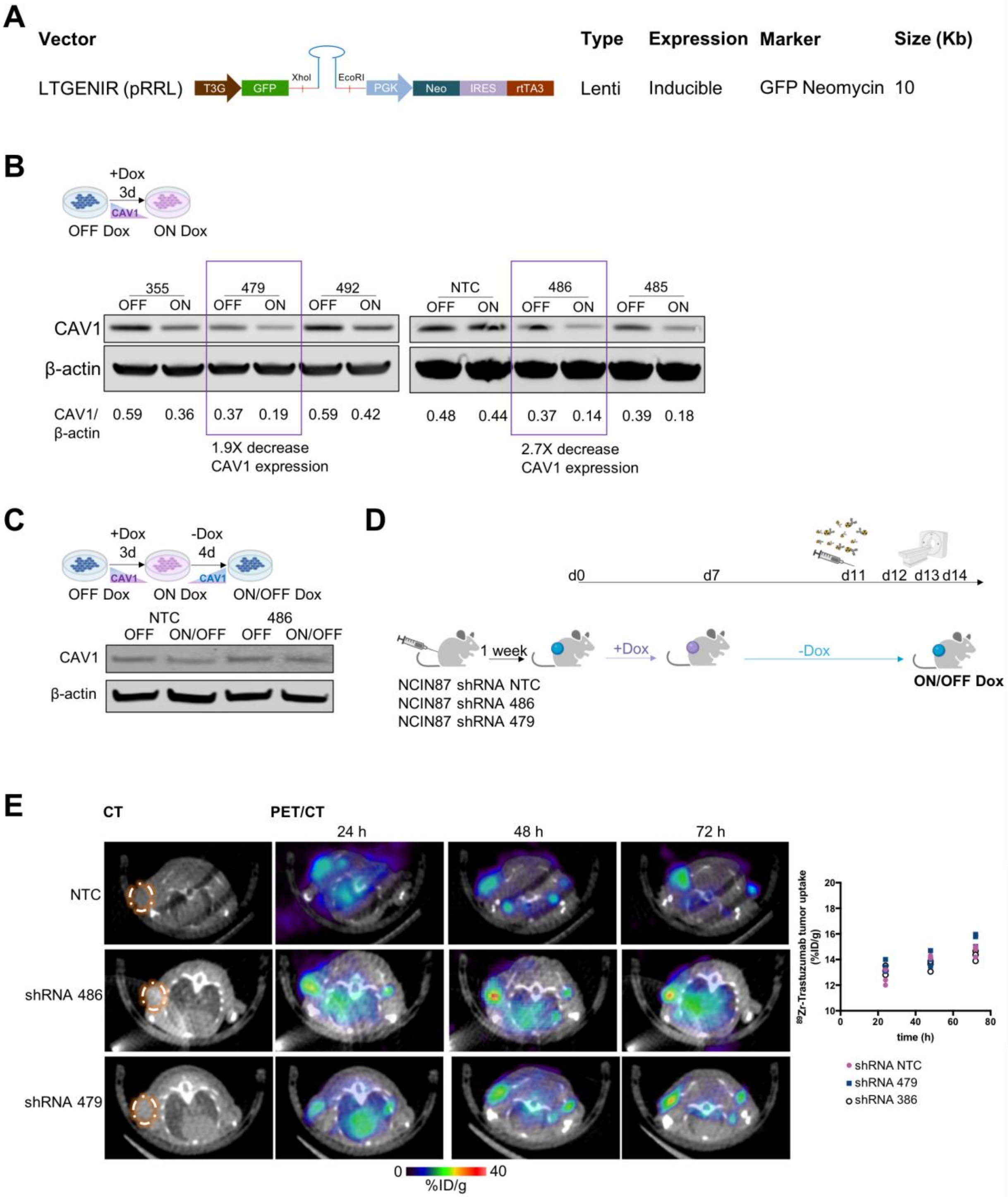


**Supplementary Figure 5.** (**A**) LT3GENIR4(pRRL) vector used to generate the tetracycline-inducible shRNA CAV1 Tet-On system. This backbone contains a neomycin selection and an inducible Dox system (Tet-On) that allows for the depletion of CAV1 in the presence of Dox. (**B**) Western blot of CAV1 in NCIN87 Tet-On systems using 5 different shRNA against CAV1 and 1 control NTC shRNA. Western blots show CAV1 depletion in cells incubated with 1 µg/mL Dox for 3 days. Dox treatment induced higher CAV1 depletion in 479 shRNA and 486 shRNA compared with 355, 492, 485 and were, therefore, used in subsequent assays. (**C**) Western blot of CAV1 in the NCIN87 Tet-On system using the shRNA 486 in the absence of Dox (OFF) or in cells treated with 1 µg/mL Dox for 3 days and then incubated with media in the absence of Dox for 4 days (ON/OFF). The ON/OFF Dox schedule allows for temporal modulation of CAV1 protein levels. (**D**) Schematic of preclinical PET imaging studies using the NCIN87 Tet-On system and an ON/OFF

Dox treatment. Athymic nude mice bearing s.c. NCIN87 shRNA NTC, shRNA 486, or shRNA 479 xenografts were orally administered with 10 mg/mL of Dox for 7 days. Mice were then orally administered PBS for 4 days. On day 11, mice were intravenously administered with [$^{89}$Zr]Zr-DFO-TDM1 (6.66–7.4 Mbq, 45–50 µg protein) and PET images recorded at 24, 48, and 72 h p.i. [$^{89}$Zr]Zr-DFO-TDM1. (**E**) Representative transversal PET images at 24, 48, and 72 h p.i. [$^{89}$Zr]Zr-DFO-TDM1 in athymic nude mice bearing s.c. NCIN87 Tet-On xenografts and ON/OFF Dox. Xenografts were obtained using NCIN87 shRNA NTC, shRNA 479, and shRNA 386 cell lines. Dox treatment and $^{89}$Zr-labeled TDM1 injection were performed as described in panel **D**. The graph plots percentage of injected dose per gram (%ID/g) of TDM1 in tumors, calculated by quantifying regions of interest (ROIs) in the PET images.

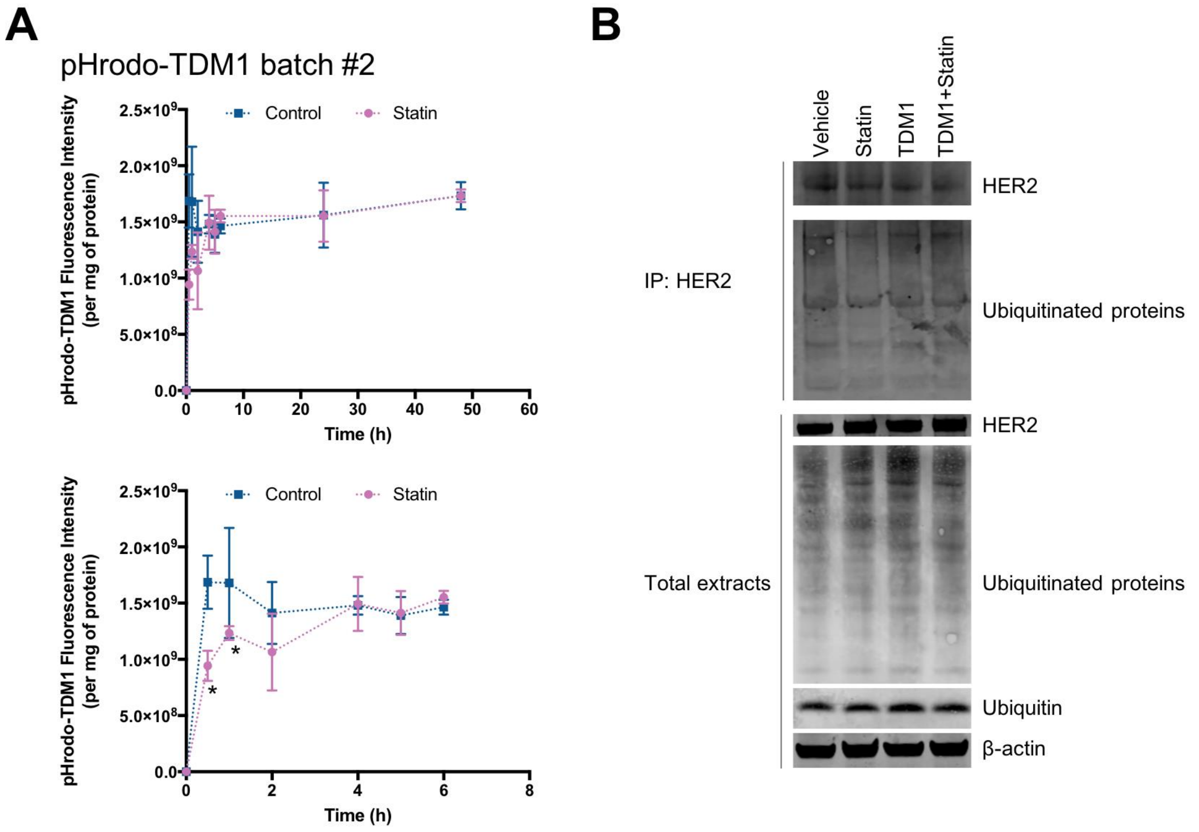


**Supplementary Figure 6.** (**A**) TDM1 internalization in HER2-expressing CAV1-positive NCIN87 gastric cancer cells in the presence and absence of lovastatin. TDM1 was conjugated to a red fluorescent pH-sensitive dye (pHrodo-TDM) in two different experimental batches (experiments obtained with Batch #1 are shown in **Figure 3B**). NCIN87 cancer cells were incubated with 1 µg/mL of pHrodoTDM1 for 30 min at 4°C. Cells were then released at 37°C, and the fluorescent signal was quantified at 30 min, 1 h, 2 h, 4 h, 5 h, 24 h, and 48 h hours after incubation with pHrodo-TDM1. The pHrodo-TDM1 fluorescent signal was normalized to the amount of protein at each time-point (*$P < 0.05$ based on a Student's *t*-test, $n = 4$). (**B**) Western blots are shown for HER2, ubiquitinated proteins, and ubiquitin from total NCIN87 cells extracts or extracts obtained after immunoprecipitation (IP) with anti-HER2 antibodies. Cells were incubated with TDM1 10 µg/mL of TDM1 or vehicle as control, together with lovastatin in the presence of the proteasome inhibitor MG-132 (10 µM) for 4 h at 37°C. β-actin was used as the loading control.

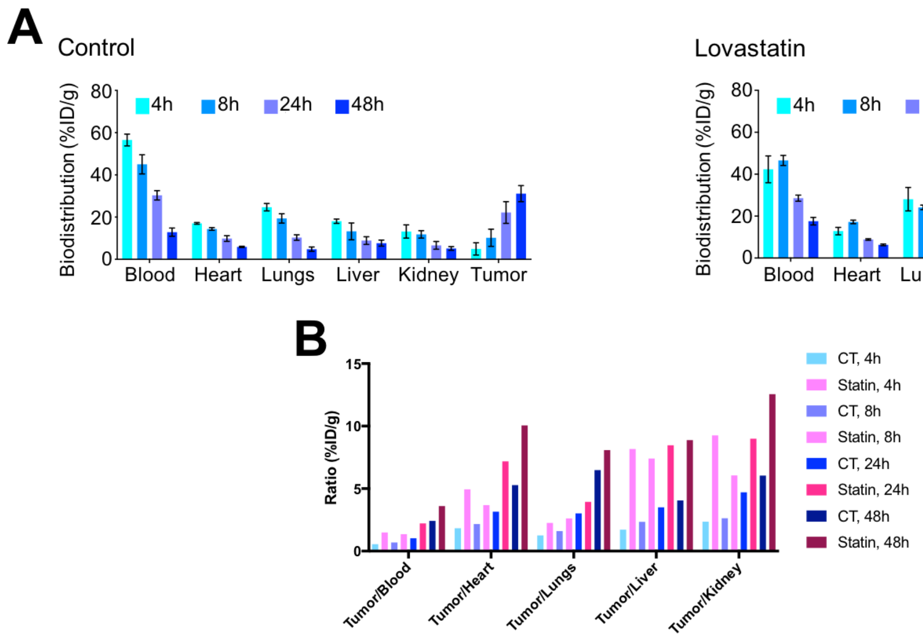


**Supplementary Figure 7.** Biodistribution data (**A**) and tumor-to-organs ratio (**B**) at 4, 8, 24, and 48 h p.i. of [$^{89}$Zr]Zr-DFO-TDM1 in athymic nude mice bearing s.c. NCIN87 gastric tumors. Lovastatin (8.3 mg/kg of mice) was orally administrated 12 h prior and at the same time as the tail vein injection of [$^{89}$Zr]Zr-DFO-TDM1 (6.66–7.4 Mbq, 45–50 µg protein). Bars, $n = 4$ mice per group, mean ± S.E.M. %ID/g, percentage of injected dose per gram.

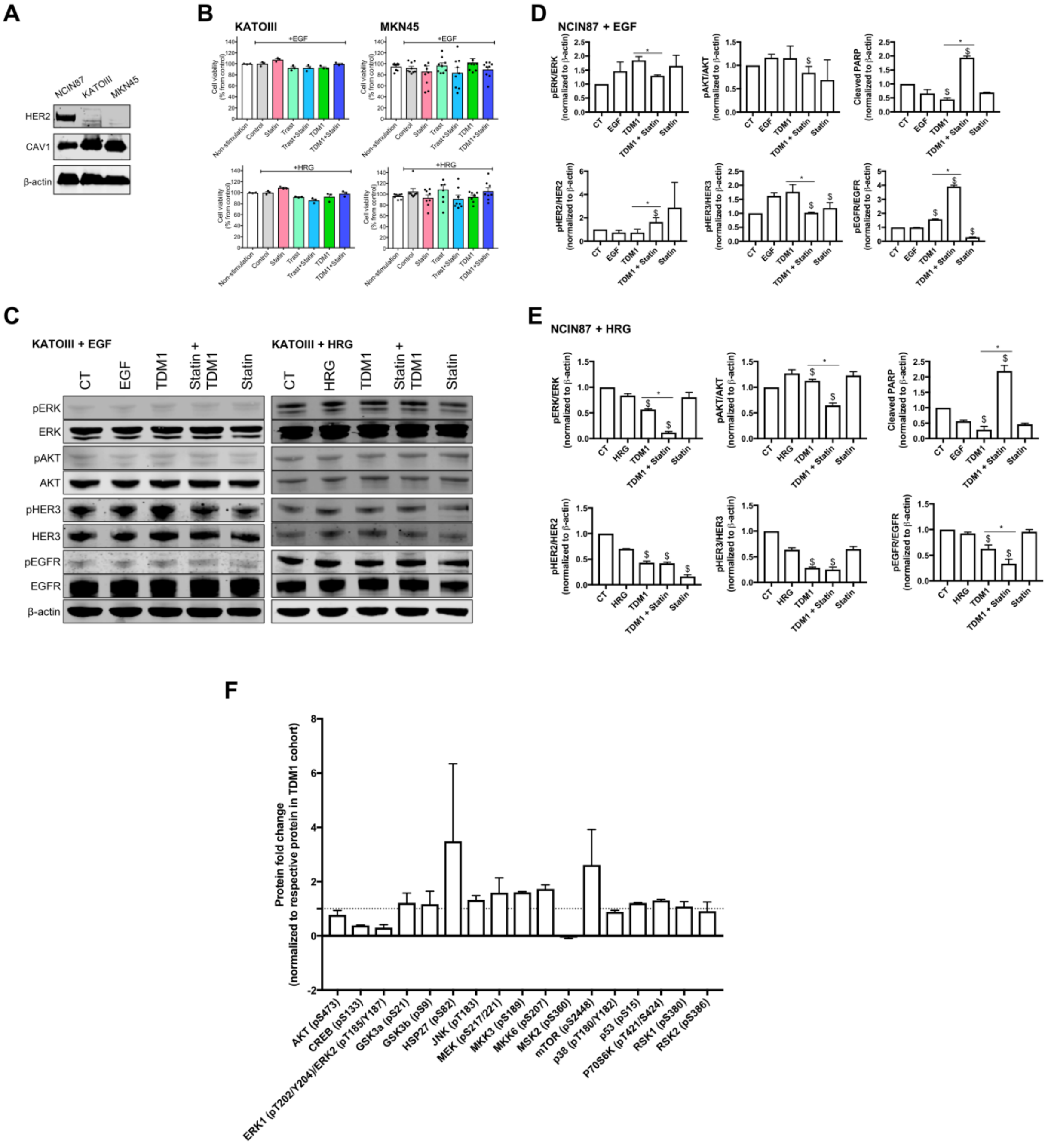


**Supplementary Figure 8.** (**A**) Western blot of HER2 and CAV1 in the total lysates of NCIN87, KATOIII, and MKN45 gastric cancer cells. (**B**) Cell viability of HER2-negative KATOIII and MKN45 gastric cancer cells at 48 h after cells incubation with trastuzumab and TDM1 alone or in combination with lovastatin. Cancer cells stimulated with 100 ng/mL EGF or HRG were incubated with 20 nM trastuzumab or TDM1. Lovastatin was added at a concentration of 25 μM. Non-stimulated cells were incubated in media in the absence of antibody or growth factor-treated samples. Bars, $n \geq 3$ per group, mean ± S.E.M. (**C**) Western blots of HER2-negative KATOIII GC cells at 48 h after cells incubation with TDM1 alone or in combination with lovastatin. (**D,E**) Quantification of western blots of HER2 signaling of HER2-expressing/CAV1$^{HIGH}$ NCIN87 gastric cancer cells stimulated with EGF (**D**) or HRG (**E**) and treated with TDM1 alone or in combination lovastatin. Bars, $n \geq 3$ per group, mean ± S.E.M. $^{\$}P < 0.05$ compared with control, $^{*}P < 0.05$ compared with TDM1.

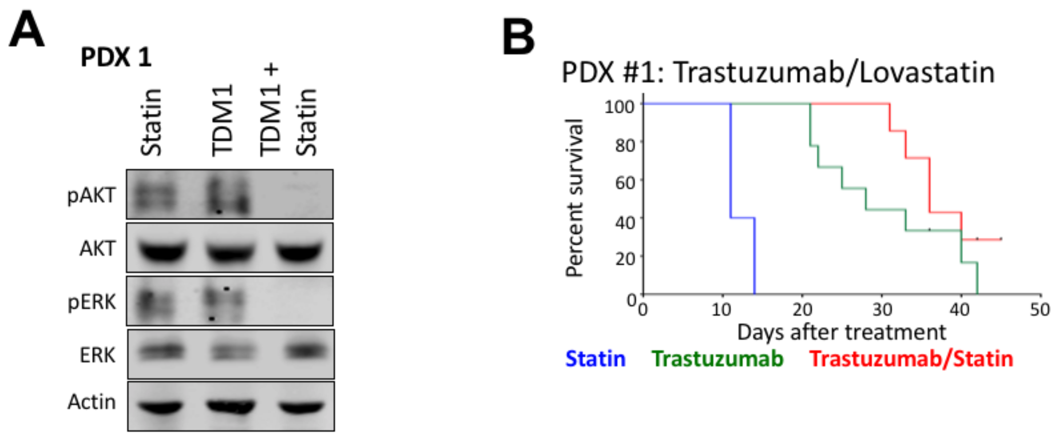


**Supplementary Figure 9.** (**A**) Western blot analyses of AKT and ERK protein expression and phosphorylation in HER2$^{+}$/CAV1$^{HIGH}$ gastric PDX #1 at 35 days after treatment with lovastatin, TDM1, or TDM1/lovastatin. Intravenous TDM1 administration 5 mg/kg weekly (for 5 weeks) was started at day 0. Lovastatin (4.15 mg/kg of mice) was orally administrated 12 h prior to and simultaneously with the intravenous injection of TDM1. (**A**) Kaplan-Meier survival curves of NSG mice bearing subcutaneous HER2$^{+}$/CAV1$^{HIGH}$ gastric PDX #1 treated with lovastatin, trastuzumab, or a combination or trastuzumab/lovastatin ($n \geq 8$ mice per group). Intraperitoneal trastuzumab administration 5 mg/kg weekly (for 5 weeks) was started at day 0. Lovastatin (4.15 mg/kg of mice) was orally administrated 12 h prior to and simultaneously with the intraperitoneal injection of trastuzumab.

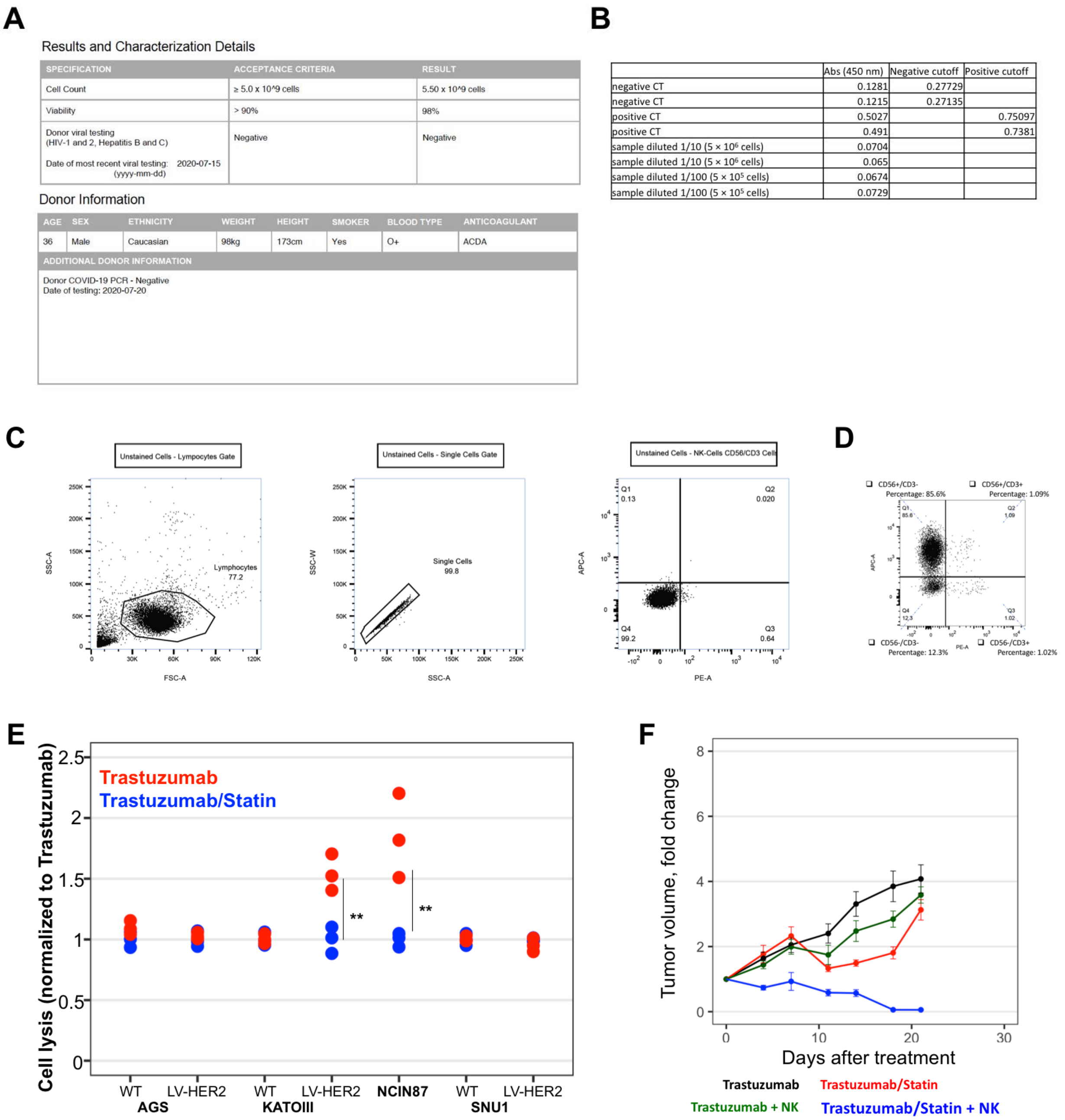

**A**

Results and Characterization Details

| SPECIFICATION | ACCEPTANCE CRITERIA | RESULT |
|---|---|---|
| Cell Count | ≥ 5.0 x 10^9 cells | 5.50 x 10^9 cells |
| Viability | > 90% | 98% |
| Donor viral testing (HIV-1 and 2, Hepatitis B and C) Date of most recent viral testing: 2020-07-15 (yyyy-mm-dd) | Negative | Negative |

Donor Information

| AGE | SEX | ETHNICITY | WEIGHT | HEIGHT | SMOKER | BLOOD TYPE | ANTICOAGULANT |
|---|---|---|---|---|---|---|---|
| 36 | Male | Caucasian | 98kg | 173cm | Yes | O+ | ACDA |

ADDITIONAL DONOR INFORMATION

Donor COVID-19 PCR - Negative
Date of testing: 2020-07-20

**B**

| | Abs (450 nm) | Negative cutoff | Positive cutoff |
|---|---|---|---|
| negative CT | 0.1281 | 0.27729 | |
| negative CT | 0.1215 | 0.27135 | |
| positive CT | 0.5027 | | 0.75097 |
| positive CT | 0.491 | | 0.7381 |
| sample diluted 1/10 (5 × 10⁶ cells) | 0.0704 | | |
| sample diluted 1/10 (5 × 10⁶ cells) | 0.065 | | |
| sample diluted 1/100 (5 × 10⁵ cells) | 0.0674 | | |
| sample diluted 1/100 (5 × 10⁵ cells) | 0.0729 | | |



**Supplementary Figure 10.** (**A**) Information on the leukapheresis pack used for NK cell isolation. (**B**) ELISA test showing negative COVID-19 result on the day of NK cells isolation using a COVID-19 IgG ELISA Kit. The negative result for COVID-19 was further confirmed on the PCR result as shown in panel **A**. (**C, D**) Gating strategy used to identify NK cells by FACs. The EasySep Human NK Cell Enrichment Kit (negative selection) was used to separate NK cells from the leukapheresis pack. The isolated cells were fluorescently stained with APC CD56 and PE CD3, and analyzed by FACs. The NK cell population was confirmed by FACs as $CD3^{-}CD56^{+}$ cells. (**E**) Ability of trastuzumab or trastuzumab/lovastatin to mediate *in vitro* ADCC as determined with NK cells from a healthy donor. GC cells were incubated for 6 h with NK cells at a ratio of 50:1 (NK:GC cells) in the presence of 100 µg/mL of trastuzumab in the absence or presence of lovastatin. Lovastatin enhances trastuzumab-mediated *in vitro* ADCC in $HER2^{+}$ CAV1-expressing cells. Points, $n = 3$, $^{**}P < 0.01$ based on a Student's *t*-test. (**F**) Therapeutic studies of NSG mice and NSG mice humanized with freshly isolated NK cells. NSG mice bearing subcutaneous $HER2^{+}/CAV1^{HIGH}$ NCIN87 xenografts were treated with lovastatin, trastuzumab, or a combination or trastuzumab/lovastatin ($n \geq 8$ mice per group). Intraperitoneal trastuzumab (5 mg/kg for 3 weeks) was started at day 0. Lovastatin (4.15 mg/kg of mice) was orally administrated 12 h prior and at the same time as the injection of trastuzumab.

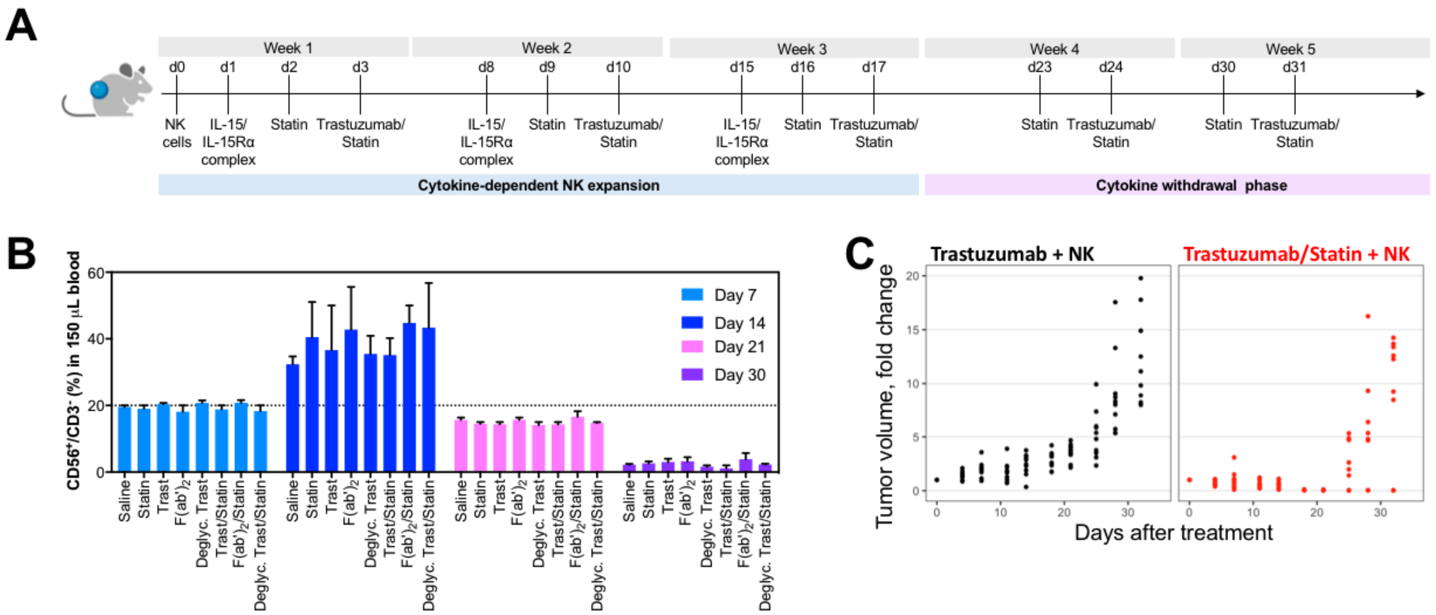


**Supplementary Figure 11.** (**A**) NSG mice bearing NCIN87 xenografts were intravenously injected with $1 \times 10^6$ human NK cells at day 0. One day after NK cells tail vein injection, the IL-15/IL-15Rα complex was intraperitoneally administered at a dose of 1.25 µg/mouse. Trastuzumab or trastuzumab/lovastatin efficacy was then evaluated during a cytokine-dependent NK expansion phase (week 1 to week 3) and a cytokine withdrawal phase (week 4 to week 5). (**B**) Percentage of human NK cells in 150 µL of murine blood collected at day 7, day 14, day 21, and day 30 after intravenous injection of NK cells. Bars, $n = 2$ mice per group, mean ± S.E.M. (**C**) Tumor volumes (fold change) as a function of time in trastuzumab or trastuzumab/lovastatin in individual NSG mice humanized with NK cells and bearing NCIN87 xenografts ($n$ = 10 mice per group). In the trastuzumab/lovastatin cohort, 3/10 mice showed remnant tumors during the NK phase expansion (week 1–3) and cytokine withdrawal phase (week 4–5). On the other hand, 7/10 mice in the trastuzumab/lovastatin cohort demonstrated similarly sized tumors compared with the trastuzumab cohort during the cytokine withdrawal phase.

A)

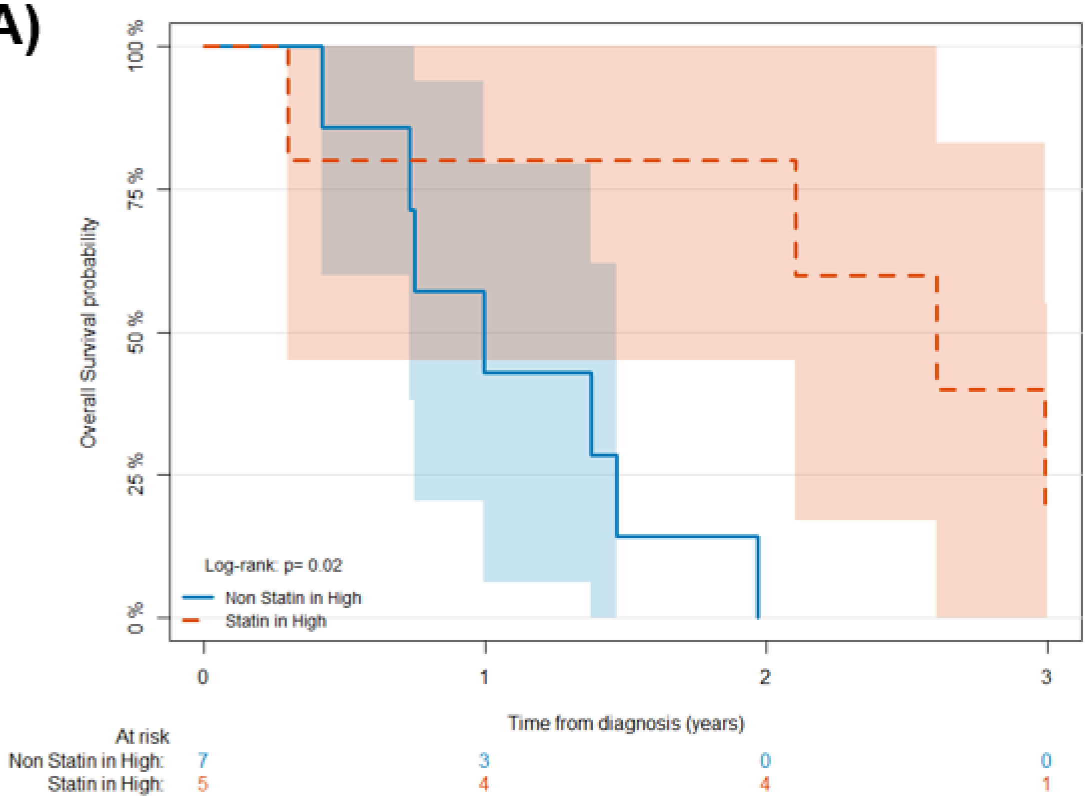


| Characteristic | At 2 years | At 3 years |
|---|---|---|
| Group | | |
| Non-Statin in CAV1-high | 0% | 0% |
| Statin in CAV1-high | 80% (45%, 100%) | 20% (0%, 55%) |

**Supplementary Figure 12.** Kaplan-Meier analysis of statin use and HER2-expressing CAV1-high GC disease outcome in patients treated with trastuzumab. CAV1-high HER2$^+$ GC without statin treatment (blue color, $n$ = 7) have a worse survival than patients treated with statin (red color, $n$ = 5). *Log rank; p=0.02*.
